# Characterizing the impacts of exotic species on the morphology of solitary threespine stickleback (*Gasterosteus aculeatus*) populations in southwestern British Columbia

**DOI:** 10.1101/2024.10.25.620354

**Authors:** Hannah M. Kienzle, Steven M. Vamosi

## Abstract

Exotic species are one of the greatest threats to native species, communities, and ecosystems. Introductions of multiple exotic species into an environment may have different effects on native populations compared to when exotic species are introduced individually. Threespine stickleback species pairs (*Gasterosteus aculeatus*) and neighbouring solitary populations in southwestern British Columbia, a textbook example of an adaptive radiation, are now under threat from multiple exotic species. We assessed whether variation in morphological characters and body shape among solitary threespine stickleback populations was associated with different combinations of introduced smallmouth bass (*Micropterus dolomieu*) and signal crayfish (*Pacifastacus leniusculus*). We also examined morphological changes over time spans of 18-43 years to determine whether contemporary characteristics have responded to the presence of exotic species. We found clear differences in stickleback traits and body shape among exotic species combinations. Stickleback coexisting with bass and crayfish were highly armoured, whereas bass-only lakes contained stickleback with reduced armour. Two stickleback populations that coexist with signal crayfish showed significant increases in size over time. These patterns suggest that smallmouth bass and signal crayfish may have significant and different impacts on stickleback morphology.

## 1. Introduction

Exotic species have been identified as one of the primary threats to the persistence of native biodiversity [1–3]. While much attention is understandably paid to extirpation and extinction [4–7], species introductions may also lead to evolution in native species. These impacts are typically harder to detect and document due to a lack of long-term survey data, although there are some examples. A gradual increase in body size and reduction of gape size has been documented in two native snakes preying on invasive and toxic cane toads over an 80-year period following their introduction to Australia, presumably as an evolutionary response to reduce and dilute toxins [8]. Similar patterns have also been observed with exotic predators and native prey. For example, dog whelk has been found to have thicker shells following exposure to exotic green crabs [9]. Invaders can drive gradual changes in native populations by impacting their range, population size, and ability to reproduce. Thus, an exotic species may not eliminate a native population, but instead reduce their numbers enough to lead to severe bottlenecks and major reductions in genetic diversity in remaining individuals [10,11]. Over longer periods of time, natural selection on native species being preyed on by an exotic predator may result in evolutionary responses that increase the likelihood of their long-term persistence [12].

Threespine stickleback (*Gasterosteus aculeatus*) display a considerable degree of morphological diversity, which presumably reflects adaptation to different environmental conditions and biological communities [13]. The range of threespine stickleback spans across the northern hemisphere in both marine and freshwater bodies, where colonization by freshwater stickleback occurred following glaciation [14]. The freshwater ‘form’ tends to show greater among-population morphological variation [14]. On the west coast of North America, freshwater populations of threespine stickleback are found primarily in coastal areas and range from Alaska to Baja California [14,15]. In these regions, there is a large degree of divergence in freshwater threespine stickleback morphology among lakes that has been attributed to environmental variation. Lake size, relative littoral area, and predator community may impact a variety of characteristics, including body shape and traits associated with foraging, predator evasion, and defense [16–18]. Populations that become isolated in freshwater bodies may evolve after only a few generations [19,20]. In addition to among-lake variation, there are cases of significant within-lake variation. Sexual dimorphism has been demonstrated in several populations, in which males consistently had larger heads and mouths relative to females [21]. A notable example of sympatric phenotypic variation occurs in several lakes that contain two distinct stickleback populations rather than one solitary population, and are referred to as benthic and limnetic stickleback [22–24].

There is a large and growing literature on the impacts of native predators on stickleback morphology. Solitary populations have been observed to coexist with a variety of native piscivores, including invertebrates, birds, and large fish [18,25,26]. Conversely, the only native fish piscivores in all species pair lakes are cutthroat trout (*Oncorhynchus clarkii*) [25]. The total quantity of piscivorous predators in a lake has been positively correlated with defensive traits such as dorsal and pelvic spine length [27]. Additionally, the presence of any native predatory fish is associated with stickleback that have a slender body shape suited for evasion that significantly differs from lakes where piscivores are completely absent [17]. When predators are common in a lake, stickleback tend to have significantly more lateral plates and longer spines [16]. Finally, there is evidence of a reduced pelvic girdle in the absence of native predators, with the degree of reduction being dependent on calcium concentrations [28–30].

Research on species pairs suggests the potential for significant negative consequences following the introduction of exotic species. Between 1988 and 1992, the species pair in Hadley Lake went extinct after the introduction of brown bullhead (*Ameiurus nebulosus*) [31]. In Enos Lake, the species pair hybridized into a single benthic-like phenotype following the introduction of signal crayfish (*Pacifastacus leniusculus*) [32]. Subsequent evidence suggested that crayfish primarily alter the reproductive behaviour of limnetic males [33]. Furthermore, it is hypothesized that signal crayfish were able to disrupt visibility and reproductive behaviour of stickleback by destroying vegetation and increasing turbidity, effectively reducing the niche for the limnetic form [32]. Thus, there is strong evidence implicating invasive species in the extinctions and phenotypic shifts of threespine stickleback species pairs. However, it is presently unclear how these outcomes can be extended to the majority of lakes, which contain a single interbreeding population of threespine stickleback.

In southwestern British Columbia, solitary stickleback populations now find themselves subject to several exotic species, with two of significant concern: smallmouth bass (*Micropterus dolomieu*) and signal crayfish. Smallmouth bass (*Micropterus dolomieu*) were introduced to Vancouver Island in the 1950’s, originally stocked legally by the government and subsequently spread illegally by anglers [34,35]. Smallmouth bass are known to significantly reduce prey fish abundance in other areas where they have been introduced, primarily through direct predation on native species [36]. In Ontario lakes, they have been associated with the extirpation of other smaller fish species similar to threespine stickleback, including brook stickleback (*Culaea inconstans*), fathead minnow (*Pimephales promelas*), and pearl dace (*Margariscus* sp.) [37]. Signal crayfish are native to mainland British Columbia and are suspected to have been introduced to Vancouver Island by anglers in the early 1900s [38]. They are recognized as having high potential for dispersal through aquatic systems and have had an impact on native crayfish populations following their introduction and spread throughout Europe [39]. Signal crayfish effects on native fish are diverse, including predation on smaller fish and eggs, competition for shelter, and destruction of vegetation, with the latter affecting the survival of juvenile fish that depend on this habitat [40,41].

Currently, we know very little of the effects of either species on solitary stickleback populations and, to our knowledge, the collective impacts of smallmouth bass and signal crayfish on stickleback have not been investigated. Previous work has identified that johnny darter (*Etheostoma nigrum*) show increased activity and vulnerability to smallmouth bass when rusty crayfish (*Oronectes rusticus*) are also present, compared to when either species was acting alone [42]. Conversely, the effects may not be additive because smallmouth bass also prey on signal crayfish, and there is evidence of stickleback comprising a larger portion of the bass’ diet in lakes with fewer crayfish present [43]. Regardless of the interaction, there is a strong possibility that exotic species in these lakes may impact stickleback morphology.

Here, we examine the potential impacts of these two exotic species on native threespine stickleback on Vancouver Island and along the Sunshine Coast, British Columbia. Our aims were to determine whether: (1) stickleback from lakes with different combinations of signal crayfish and smallmouth bass differ significantly in their morphology from those in lakes without exotic species, and (2) any significant morphological changes have occurred over time in lakes containing smallmouth bass and signal crayfish, by comparing contemporary fish to historical samples. In smallmouth bass lakes, we predict that contemporary stickleback would be more limnetic-like with increased armour, due to a greater risk of bass predation. In signal crayfish lakes, we predict a more benthic-like form, based to the outcome of the hybridization of the Enos Lake species pair where crayfish were implicated in the impairment of reproductive limnetic males. In lakes with both species present, we predicted fish would resemble stickleback from lakes without exotic species, because of smallmouth bass predation on signal crayfish.

## 2. Methods

### 2.1. Study area

Ten coastal lakes known to contain threespine stickleback on southern Vancouver Island, Gulf Islands, and along the Sunshine Coast were selected for surveys in spring 2016 (Fig. 1, Table 1). All lakes are in separate watersheds with the exception of Blackburn and Cusheon Lake. These lakes were specifically chosen because they possess similar abiotic characteristics (Table 1) and native fish compositions and a range of different non-native species combinations (Table 2).

**Figure 1.**
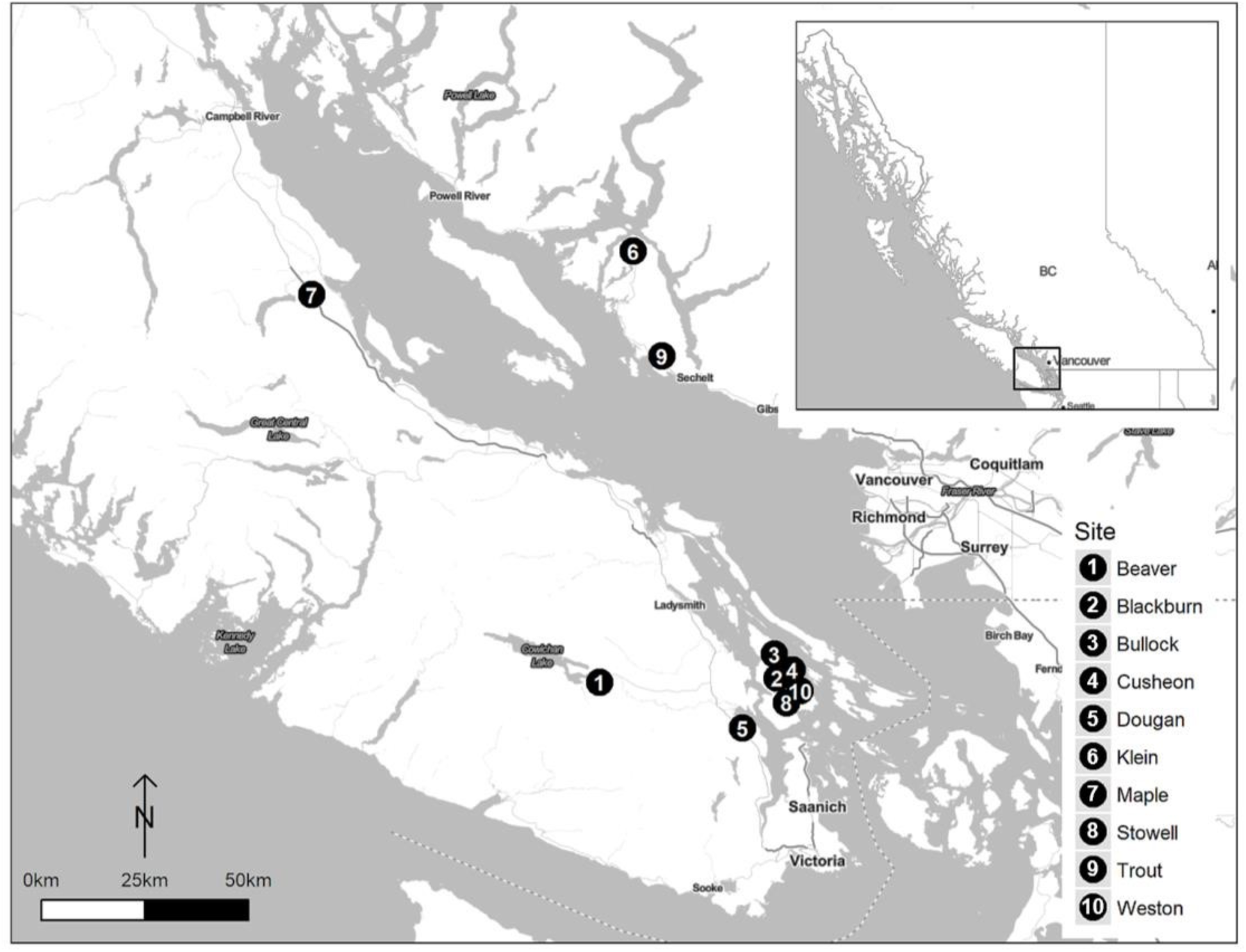
A map of the south coast of Vancouver Island and Sunshine Coast, with the locations of 10 lakes that were sampled in 2016. Inset map shows the region relative to the province of British Colombia, Canada.

**Table 1.** Locations and environmental characteristics of 10 lakes sampled on Vancouver Island, Gulf Islands, and Sunshine Coast.

| Lake | Abbreviations | Coordinates | Surface area (Ha) | Max depth (m) | Perimeter (km) | Elevation (m) |
| --- | --- | --- | --- | --- | --- | --- |
| Beaver | BV | 48°48'46.68"N<br>124° 4'44.34"W | 18 | 7.5 | 1.8 | 183 |
| Cusheon | CU | 48°48'59.90"N<br>123°28'10.62"W | 28 | 9.1 | 3.9 | 99 |
| Bullock | BU | 48°52'24.58"N<br>123°30'31.49"W | 10 | 6.9 | 1.3 | 33 |
| Blackburn | BB | 48°49'21.26"N<br>123°29'5.56"W | 4 | 7.5 | 0.8 | 106 |
| Klein | KL | 49°43'48.56"N<br>123°58'9.01"W | 35 | 36 | 2.7 | 142 |
| Maple | MP | 49°38'17.82"N<br>125° 1'2.08"W | 20 | 12.8 | 2.5 | 136 |
| Dougan | DG | 48°42'52.09"N<br>123°36'49.91"W | 10 | 22 | 1.5 | 45 |
| Weston | WE | 48°47'3.35"N<br>123°25'28.03"W | 18 | 12.2 | 2.1 | 69 |
| Trout | TR | 49°30'29.69"N<br>123°52'34.46"W | 7 | 17.4 | 1.3 | 157 |
| Stowell | SW | 48°46'54.15"N<br>123°26'38.86"W | 6 | 7.5 | 1.0 | 77 |

**Table 2.** Summary of predator composition and stickleback samples from the 10 studied lakes. Stickleback sample size is indicated by the total, followed by (M/F) breakdown in brackets. Historical data samples sizes are listed in the format of dataset A total (M/F)/dataset B total. Species abbreviations: CS = coho salmon, CT = cutthroat trout, PK = Pumpkinseed, PS = Prickly Sculpin, RB = rainbow trout, SC = signal crayfish, SMB = smallmouth bass.

| Lake | Contemporary sample size | Historical sample size | SMB | SC | Other fish species | Historical and contemporary notes |
| --- | --- | --- | --- | --- | --- | --- |
| Beaver | 39 (25/14) | 40 (16/24) / 41 | + | + | CT, RB, CS, PS | SMB present, but very few. None seen or reported from 1983 to 2002. Recommendation in 2002 to reduce stickleback population to boost CT. No PS reported but were caught during 2016 surveys. |
| Cusheon | 18 (5/13) | — | + | + | CT, RB | SMB reported in 2001, 2004, 2005. Schools of SMB fry observed during 2016 surveys. HabitatWizard reports PS in 1995, but none caught during four days of surveys in 2016. |
| Bullock | 40 (36/4) | 30 (15/15) | + | — | CT, RB | Used as a fish farm from 1967-1979. No fish except stickleback reported in 1981, extremely eutrophic. No SMB in 2003 but reported in 2009. |
| Blackburn | 35 (5/30) | — | + | — | CT, RB | SMB absent from 1972 reports but spotted in 2001. |
| Klein | 35 (15/20) | 40 (22/18) / 25 | — | + | CT, RB | No stickleback found in CT stomachs in 1988, discussion of stickleback transplants from other lakes. No signal crayfish reported in 1988, but reports of crayfish in surrounding lakes in the same watershed in 2005 (Bondar 2005). |
| Maple | 25 (19/6) | — | — | + | CT, RB | Signal crayfish absent from surveys in 2011-2012 (Beck 2013). |
| Dougan | 31 (9/22) | 27 (10/17) / 38 | — | + | CT, RB, PK, CS | Pumpkinseed first reported in 2002. Survey identified crayfish (*Potamobius sp.*) in 1951. |
| Weston | 34 (19/15) | — | — | — | CT, RB |  |
| Trout | 30 (18/12) | 26 (13/13) / 37 | — | — | CT | Only CT in all reports. |
| Stowell | 30 (13/17) | — | — | — | CT, RB | No stickleback in CT stomachs in 1984. Stocked with CT up to 2006. |

They can be collectively described as relatively small, shallow, and having similar native predators, with most containing only cutthroat and rainbow trout (*Oncorhynchus clarkii, Oncorhynchus mykiss*) and occasionally coho salmon (*Oncorhynchus kisutch*). One notable exception is Dougan Lake, where non-native pumpkinseed sunfish (*Lepomis gibbosus*) were more abundant in traps than stickleback. Additionally, although not noted in previous databases, Beaver Lake was found to contain prickly sculpin (*Cottus asper*), which have been identified as having a significant influence over threespine stickleback morphology [44].

Signal crayfish presence was confirmed in 2016 for the five lakes they had been previously reported in, as they were caught in minnow traps during stickleback sampling. Lake characteristics and presence of native and non-native predators were determined using lake reports accessible through HabitatWizard (https://maps.gov.bc.ca/ess/hm/habwiz/).

### 2.2. Sample collection

Gee minnow traps were placed in the littoral zone at depths of ≤5m for a maximum of 24 hr. A total of 317 threespine stickleback were caught from May 8^th^ to June 15^th^ of 2016 (i.e., during the breeding season when adult fish are predominantly inshore). Only adult stickleback with a standard length ≥30mm were kept, with smaller stickleback and all bycatch being immediately released. Fish were euthanized on-site via immersion in a solution of eugenol (clove oil) at a concentration greater than 400mg/mL. All procedures were carried out in accordance with Canadian Council of Animal Care standards (University of Calgary Animal Care Protocol AC15-0033). Fish were immediately preserved in 10% neutral buffered formalin, with dorsal and caudal fins clipped and stored separately in 95% ethanol to be used for genetic differentiation of males from females. To accurately measure bony structures, fish were stained with an alizarin red solution following a standard procedure and placed in 40% isopropanol for storage.

In addition to samples from 2016, threespine stickleback captured from 1973 to 1982 from Beaver, Dougan, Klein, and Trout lakes were obtained from the Beaty Biodiversity Museum, University of British Colombia. Differences in preservation techniques can have a significant effect on morphology [26], but because historical and contemporary samples were fixed in formalin, stained, and stored using similar procedures, any shape variation due to preservation is likely negligible. In addition, four months elapsed between fixation in formalin and preservation in isopropanol and measurement of traits, which reduces the likelihood of size disparity between contemporary and museum fish introduced by shrinkage of fish over time [45]. Finally, the likely negligible effects of distortion are expected to be random with respect to the subsequent presence/absence of exotic species in lakes.

Historical and contemporary individuals were photographed in the same manner using a Nikon Coolpix P530, on both the left lateral side and the ventral side. Camera set up and positioning of each fish were standardized in all photos to minimize parallax.

### 2.3. Trait measurements

Stickleback were measured using *ImageJ* software version 1.48 [46]. A total of 15 traits were determined from photos: standard length, eye diameter, gape length, snout length, head length, head depth, body depth, dorsal spine lengths, ectocoracoid length, caudle peduncle length and width, pelvic girdle length and width, and pelvic spine length (Fig. S1). In cases where measurements from photographs were not possible (due to bending, angle of spines, etc.) traits were measured using digital calipers under a dissecting scope. Two additional meristic traits, number of lateral plates and number of gill rakers on lower arch, were counted under a dissecting scope. Lateral plates were counted from the left side of each fish, whereas gill rakers were dissected and counted from the right side to avoid obscuring other traits, in the event that they needed to be re-measured at a later date. To visually assess effective cross-sectional size, which can limit ingestion by predatory fish, we estimated average diameter for each lake and across time. Population means for body depth, second dorsal spine length, pelvic girdle width, and pelvic spine length were used to calculate maximum cross-section size with full extension of the spines (see Supplementary Methods for details).

### 2.4. Body shape

Geometric morphometrics allow for body shape to be characterized and quantitatively described using a set of biologically relevant landmarks on each individual. Twenty-five landmarks were applied to 315 photos using *tpsDig* v. 2.30 [47] and *tpsUtil* v. 1.74 [48]. Only landmarks that were homologous and clearly identifiable on every fish were selected, following most of the landmarks delineated by [49] (Fig. S1). To determine accuracy of placement, landmarks from a random subset of individuals were placed a second time; the shape of these replicate fish did not differ significantly from the first set (F_1,18_ = 0.267, p = 0.973).

### 2.5. Sex determination

The sex of each fish was determined using a sex specific polymorphism on the *Idh* gene. Molecular methods were used instead of dissection in order to maintain body structure for other analyses. Furthermore, genetic identification using the *Idh* gene has been found to have 99% accuracy in matching to phenotypic sex [50]. DNA from fin clips was extracted using a QIAGEN DNeasy Blood and Tissue Kit (QIAGEN, Valencia, CA). Extracted DNA was amplified using a published PCR protocol [51]. DNA was visualized on a 2% agarose gel, with males having 302 and 271 base pair bands, and females having only the 302 base pair band. Eighty-three gravid females were identifiable either in photographs or under a dissecting scope and did not need to be sexed using molecular methods. Ten individuals were used to verify the identity of the other cryptic stickleback – all gravid females from the subset had a single 302 base pair band and were compared to the rest of the samples in each run of PCR.

### 2.6. Trait analyses

All linear measurements were ln(x+1) transformed and examined for outliers. Although body size can potentially influence trait size, traits were not adjusted according to the standard length of each individual. Allometric effects in specific traits of threespine stickleback can differ among environments [52] and only a small amount of variation in shape is typically explained by size [53]. Several of the traits (spine lengths, lateral plates, and gill rakers) were completely uncorrelated with size whereas others were correlated only in some lakes. In addition, correcting for size consumes four of the trait measurements (standard length, body depth, head depth, and snout length), which may contain biologically meaningful variation that could otherwise be examined and compared among sites and between time periods.

To examine differences in phenotypic traits among populations, contemporary data consisting of 17 meristic and linear trait measurements were split by sex and summarized using a Principal Component Analysis (PCA), which allows a large number of response variables to be reduced to a few uncorrelated components while still retaining most of the variation [54]. Measurements with loadings of 0.5 and above were considered to be substantial contributors to a component. PCA was completed using the *psych* package in R [55]. The main components were then analyzed with lake assemblages and non-native species combinations using a MANOVA, to assess whether there were clear distinctions among these groups. This was followed by a post-hoc Discriminant Function Analysis (DFA) using the *MASS* package in R [56]. In addition, traits summarized in PC1 were regressed on four abiotic variables (elevation, perimeter, surface area, and maximum depth) to determine whether any of the variation could be explained by major features of the surrounding environment. Finally, a separate PCA combined all contemporary linear measurements with linear measurements of the Enos Lake species pair (from [57]). Because Enos Lake fish were clear examples of the benthic and limnetic form prior to their hybridization starting in the late 1990s [58,59], this pre-hybridization species pair data can be compared to contemporary non-species pair stickleback to determine where these populations lie on a benthic-limnetic scale.

To examine different facets of temporal effects on phenotypic traits, two datasets were used. The first dataset (hereafter, dataset A) consists of stickleback generally sampled in the 1970s and 1980s (Bullock was sampled in 1999) and measured in 1999-2000 by SMV [25]. These historical data were combined with our contemporary measurements and consist of six measured traits from five lakes (Bullock, Beaver, Dougan, Klein, and Trout lakes) with males and females identified. The second dataset (hereafter, dataset B) consists solely of measurements taken by HMK in 2016 of both contemporary and historical fish. This set uses all 16 traits (excluding gill raker count) with fish from four lakes (Beaver, Dougan, Klein, and Trout lakes). Because sex determination was not possible from these museum samples, males and females were pooled in this dataset. Each dataset was summarized with a PCA and evaluated with a MANOVA, splitting both by site to examine the main effects of year and sex for dataset A, and year only for dataset B.

### 2.7. Geometric morphometric analysis

Shape analyses were conducted using the package *geomorph* v. 3.0.5 in R [60]. A PCA was performed on Procrustes transformed data, and the first component exhibited bending of the vertebral column that is characteristic of preserved fish [61]. Because this variation is not biologically relevant, residuals generated from regressions on this first component were used to remove variation introduced via bending [52]. To examine contemporary body shape, landmark data were split by sex and superimposed using General Procrustes Analysis (GPA), allowing the raw landmark data to be scaled, translated, and rotated around a central origin to reduce variation caused by positioning [62]. The data were then summarized using a PCA and assessed using a Procrustes ANOVA and DFA to examine the effects of site on body shape of both males and females. For shape differences between contemporary and historical samples, the same methods were used but the data were split by site in order to examine the effects of year. Males and females were pooled for this analysis because, as previously mentioned, sex determination was not possible for the fish obtained through the museum.

## 3. Results

### 3.1. Trait measurements – contemporary lake comparisons

A PCA on 17 linear trait measurements resulted in the first two components collectively explaining 71.2% and 66.3% of the variation for females and males, respectively (Table S1). The first principal component (PC1), which explained 50.5% of the variation for females and 41.9% for males, was primarily driven by traits corresponding to size. Most of these characteristics were strongly associated with PC1 and contributed relatively equally, including standard length, head length, head depth, body depth, eye diameter, snout length, gape length, and caudal peduncle width. Pelvic girdle width and length, caudal peduncle length, and ectocoracoid length also contributed to a lesser extent and varied between males and females. The second principal component (PC2) explained 20.7% and 24.4% of the variation for females and males, respectively, and consisted of characteristics attributed to armour (Table S1). The dorsal and pelvic spines were of relatively equal importance, while lateral plates and pelvic girdle length contributed to a lesser degree.

These first two components, which explained a majority of the variation, were evaluated among lake populations with a MANOVA. Variation in morphology was highly significant among lakes and among non-native species combinations (Table 3, Fig. 2). A follow-up DFA generated correlations among the first two principal components and two linear discriminants. Armour was identified as the more important predictor of lake origin for both males and females (Table S2). When given measurements of stickleback with no lake identifier, the DFA correctly assigned 62-63% of individuals to the correct lakes, whereas only 18-20% of randomized data were correctly assigned. For both sexes, Beaver Lake individuals were the most distinct, characterized by longer spines, wider pelvic girdles, and more lateral plates (Table S2, Fig. S2). Cusheon Lake stickleback were also fairly easily classified for females and were larger in size than fish from other lakes. Male stickleback from Bullock Lake were also easily classified with smaller armour trait values compared to other lakes. Conversely, male fish from Stowell Lake and Dougan Lake were never correctly classified and were frequently sorted as Bullock and Klein Lake stickleback respectively. For female fish, Maple Lake stickleback were often misidentified as being from Klein or Dougan Lake.

**Figure 2.**
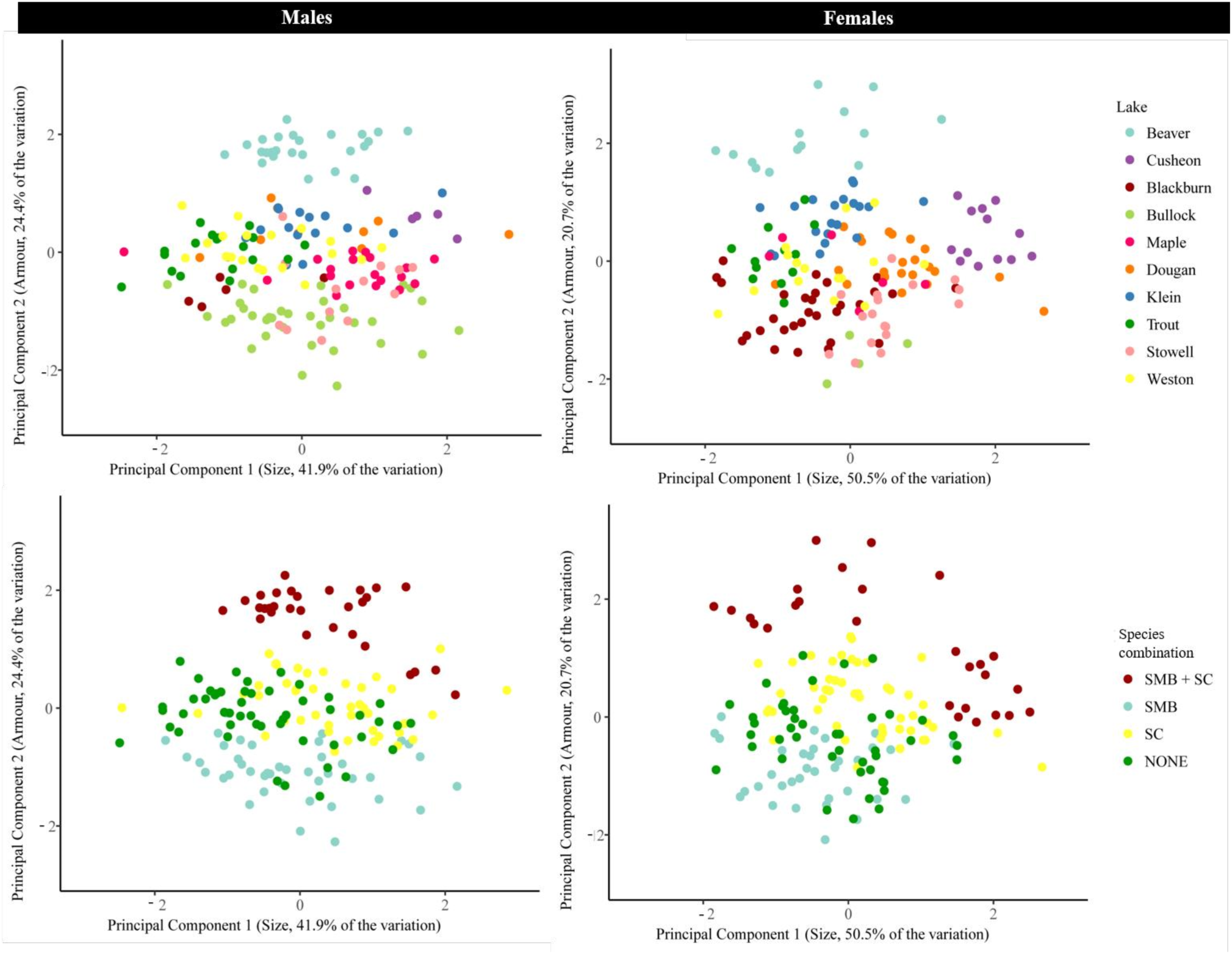
Stickleback trait variation for each lake (top row) and species combination (bottom row) for males and females. Principal Component 1 primarily represents size characteristics and Principal Component 2 primarily represents armour traits.

**Table 3.** MANOVA results for effects of lake and non-native species combination on male and female stickleback morphology, using contemporary data.

|  |  | df | Pillai | F | num df | den df | p value |
| --- | --- | --- | --- | --- | --- | --- | --- |
| Females | Lake | 9 | 1.415 | 38.448 | 18 | 286 | < 0.001 |
|  | Residuals | 143 |  |  |  |  |  |
|  | Species combination | 3 | 0.694 | 26.412 | 6 | 298 | < 0.001 |
|  | Residuals | 149 |  |  |  |  |  |
| Males | Lake | 9 | 1.248 | 28.394 | 18 | 308 | < 0.001 |
|  | Residuals | 154 |  |  |  |  |  |
|  | Species combination | 3 | 0.898 | 43.465 | 6 | 320 | < 0.001 |
|  | Residuals | 160 |  |  |  |  |  |

Armour traits were also the best predictor of exotic species combination, with 62-72% of stickleback assigned to the correct group compared to 26-33% for randomized data. Stickleback from lakes with neither non-native species had the lowest rate of correct classification (50-60%) and were often misclassified as being from lakes with just bass or just crayfish (but never a combination of both). SMB and SC lakes tended to have the longest spines, more plates, and longest pelvic girdles. Stickleback from lakes with only SMB generally had reduced/lower armour trait values, although there was some overlap with SC lakes and lakes where non-native species are absent. When examining estimates of cross-sectional diameter, which incorporate both armour and body depth, the same patterns still hold where SMB + SC lakes and SMB lakes have fish with the largest and smallest diameters, respectively (Fig. 3). There was a 36% difference in diameter between the largest (Cusheon Lake) and smallest (Bullock Lake) populations.

**Figure 3.**
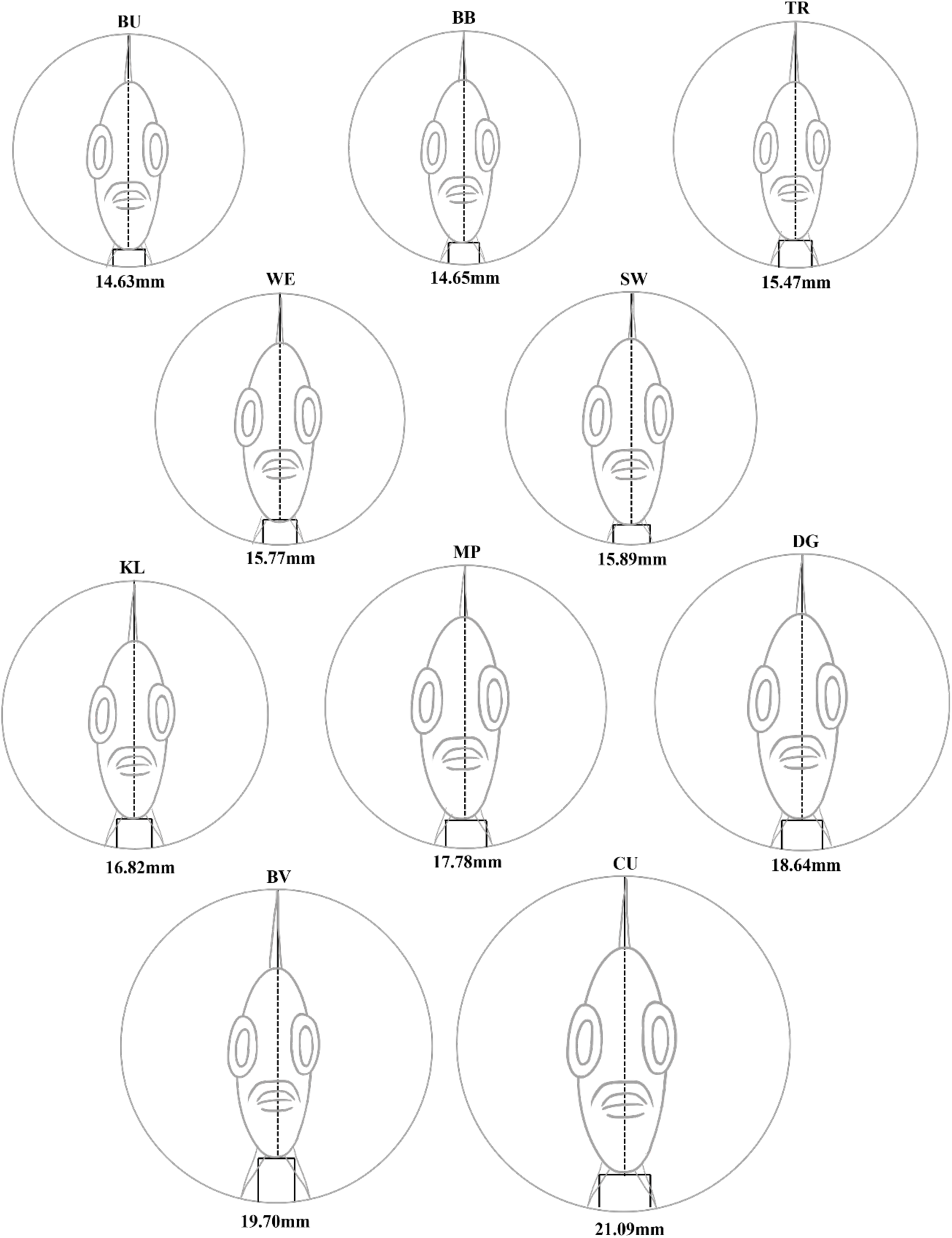
Relative estimated cross-sectional diameter of contemporary stickleback from Bullock (BU), Blackburn (BB), Trout (TR), Weston (WE), Stowell (SW), Klein (KL), Maple (MP), Dougan (DG), Beaver (BV), and Cusheon (CU) Lakes.

### 3.2. Trait measurements – temporal comparisons

For dataset A, a total of six traits were shared by the contemporary and historical data. These traits were summarized with a PCA, with the first two components explaining 86.7% of the variation (Table S4). PC1 explained 52.9% of the variation, consisting of four armour traits (three spine measurements and lateral plate count), whereas PC2 explained 33.8% of the variation, consisting of standard length and body depth. The PCA for dataset B contained more trait measurements (a total of 16) and had opposite component contributions, with PC1 consisting of size traits and PC2 consisting of armour traits, explaining 43.8% and 26.2% of the variation respectively. For both datasets, characteristics described by the first two principal components varied significantly between years for all lakes (Table S5). For dataset A, Beaver and Trout Lake differences were primarily driven by armour, whereas these differences were mostly driven by size for Klein and Bullock Lakes (Table S6). Although armour and size were nearly equally responsible for changes from past to present in Dougan Lake for dataset A, dataset B differences were primarily driven by size. Dataset A also examined the effects of sex on morphological variation, which was only significant in Dougan and Klein Lake. Dougan Lake males have more prominent armour traits but are smaller in size relative to females, whereas Klein Lake males have less armour but are larger in size compared to females. Additionally, for dataset A it is important to note that main effects could not be interpreted for Beaver Lake because there was an interaction between year and sex. However, Beaver Lake was also present in dataset B, which suggested there was a highly significant effect of lake on morphology (F_2,77_ = 82.611, Pillai’s Trace = 0.682, p < 0.001).

For both datasets, there is a clear increase in size from past to present for Klein and Dougan Lakes (Fig. S3). Additionally, Trout and Beaver Lake fish have smaller armour trait values, although this change was less pronounced in dataset A. Bullock Lake was present only in dataset A and exhibited a shift toward smaller sizes. Overall, both datasets show the same trends with the exception of Beaver Lake, which had inconsistent changes in size between datasets. Regarding stickleback diameter, the Dougan Lake population had the greatest amount of overall change and Beaver and Trout Lake populations changed the least (Table S3, Fig. S4).

### 3.3. Geometric morphometrics – contemporary lake comparisons

Bending-corrected landmarks were summarized using a PCA and examined with Procrustes ANOVA (Table 4, Table S7). Body shape varied significantly among lakes and among non-native species combinations (p = 0.01). Overall, females exhibited more variation in body shape within each lake than males (Fig. S5, Fig. S6). For males, Beaver, Klein, and Maple Lakes had fish with the ascending branch of the pelvis reaching further up the lateral side, whereas the opposite was true for Bullock Lake. Maple Lake was also characterized by a longer dorsal fin than fish from other lakes. Although female body shape is much less distinctive among lakes, Blackburn Lake individuals appear to be characterized by longer dorsal fins and shorter ascending branch of the pelvic girdle, as well as longer distance between the pelvic spine and anal fin.

**Table 4.** Principal component analysis of body shape data, for both contemporary and temporal datasets, displaying only the first five principal components. The contemporary analysis is split by males and females, whereas the temporal analysis is split by lake.

|  |  | Proportion of variance explained |  |  |  |  |  |
| --- | --- | --- | --- | --- | --- | --- | --- |
|  |  | PC1 | PC2 | PC3 | PC4 | PC5 | Cumulative |
| Contemporary analysis | Male | 0.213 | 0.123 | 0.095 | 0.013 | 0.012 | 0.456 |
|  | Female | 0.238 | 0.134 | 0.115 | 0.088 | 0.074 | 0.649 |
| Temporal analysis | Beaver | 0.403 | 0.112 | 0.089 | 0.071 | 0.056 | 0.731 |
|  | Dougan | 0.355 | 0.152 | 0.093 | 0.065 | 0.052 | 0.717 |
|  | Klein | 0.282 | 0.169 | 0.121 | 0.084 | 0.057 | 0.713 |
|  | Trout | 0.317 | 0.173 | 0.114 | 0.097 | 0.049 | 0.750 |

Specific differences between lake populations were assessed using a DFA. There was a high proportion of correct lake classifications, whereas the randomized dataset had a low proportion of individuals correctly assigned (Table S8). When evaluating body shape relative to non-native species combinations, the model classified 85-93% of individuals into the correct groups relative to the 30-36% correct classifications for a randomized dataset. For both males and females, SMB lakes had the highest proportion of correct classifications although the other combinations were highly distinct as well (Fig. S7).

### 3.4. Geometric morphometrics – temporal comparisons

Body shape varied significantly between years for fish from all four lakes (Table S7). All sites had longer anal and/or dorsal fins from 1973-1982 to 2016. Klein Lake fish had a longer ascending branch of the pelvic girdle and a slightly flatter abdomen, whereas Beaver Lake fish had a larger and rounder head shape (Fig. 4; also Fig. S8, Fig. S9).

**Figure 4.**
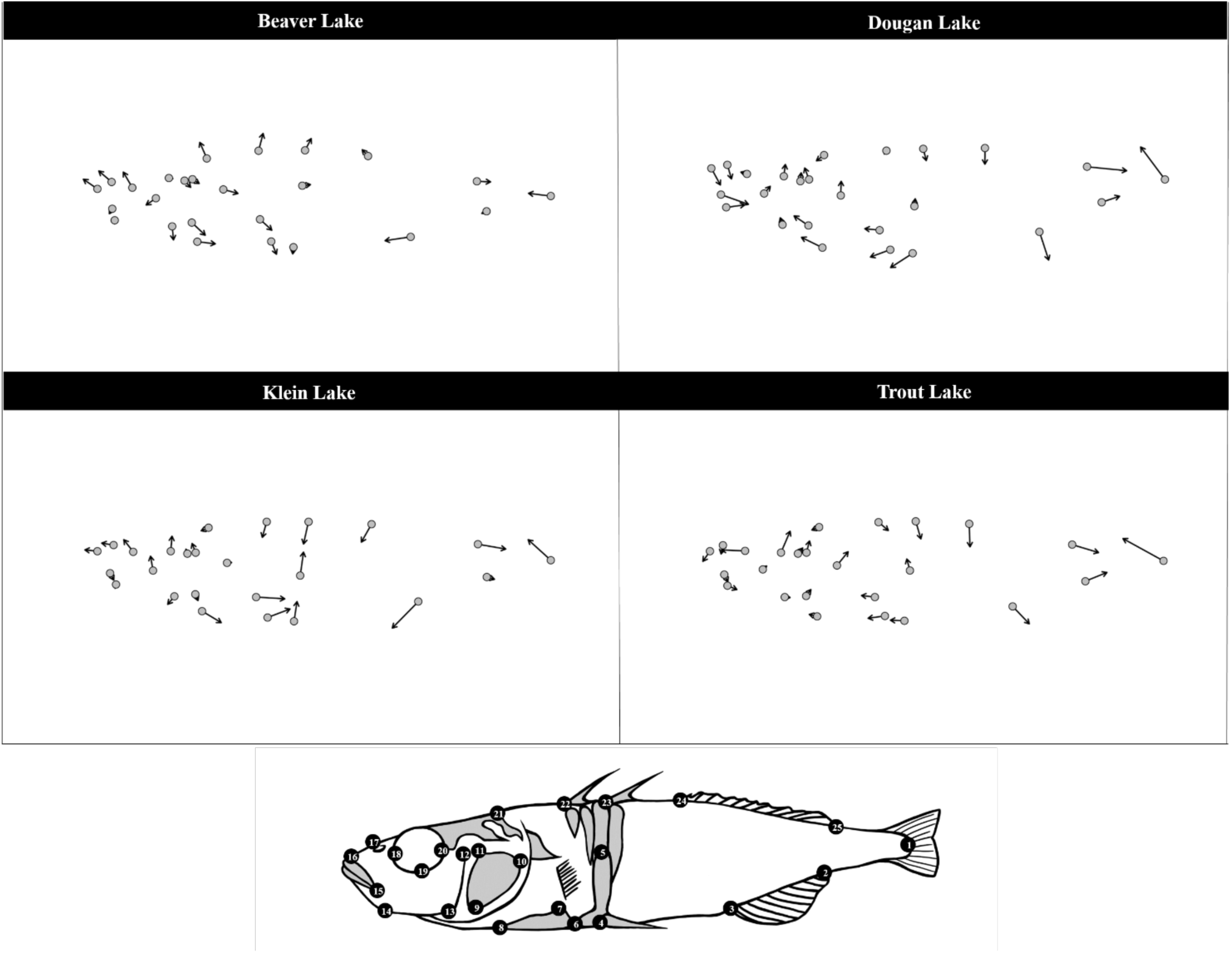
Changes in mean shape for Beaver, Dougan, Klein, and Trout Lake from 1973-1982 to 2016. Arrows indicate direction and amount of shape change over time. A landmark diagram is included for reference.

### 3.5. Comparison to Enos Lake populations

Contemporary fish generally more closely resemble the benthic phenotype than the limnetic phenotype based on the chosen armour and size traits (Fig. 5). Beaver and Cusheon Lake stickleback are most different from either species pair form, with the former having longer spines and more lateral plates and the latter being larger in body size.

**Figure 5.**
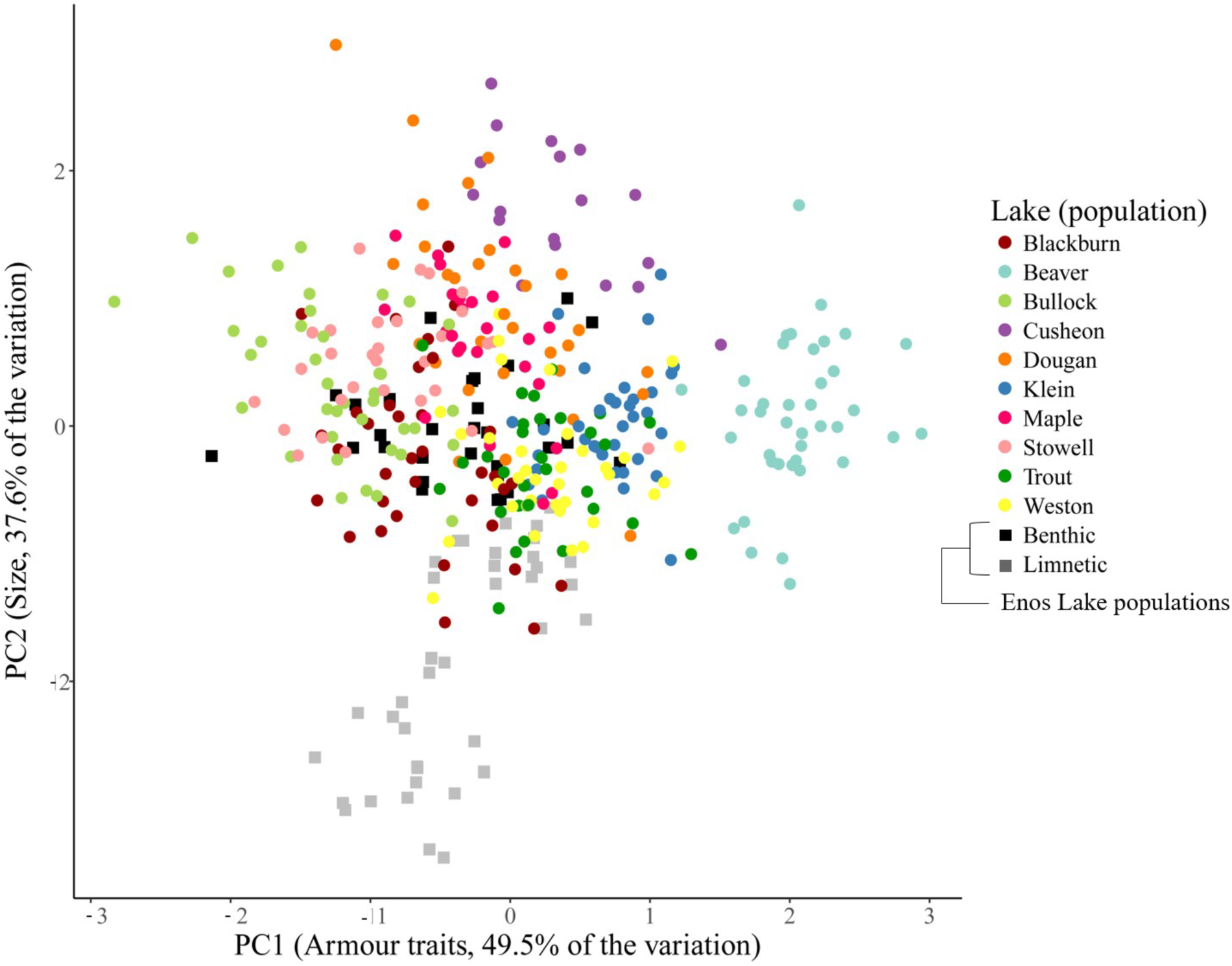
PCA of trait variation in stickleback sampled from ten lakes in 2016, with the addition of benthic and limnetic Enos Lake species pair. The first principal component represents armour traits, whereas principal component 2 consists of size-related traits.

## 4. Discussion

Stickleback morphology varied significantly among lakes and non-native species combinations. One of the more noteworthy differences among groups was between SMB + SC and SMB lakes, which had the largest and smallest armour trait values respectively, despite the fact that they both share bass presence. This suggests that the effects of non-native species could be additive, and that there might be stronger pressures from both SMB and SC, favouring greater armour.

Experimental observations indirectly support this hypothesis: SMB force johnny darters to sheltered areas and reduce their overall activity, but when rusty crayfish are added the darters are driven from shelter, their activity increases, and the prey fish are more vulnerable to both species [42]. Furthermore, signal crayfish have a similar effect on sculpin and other fish species by driving them from refuges and increasing their activity levels [63]. If stickleback are able to seek shelter in the absence of SC and avoid predation by bass, this could mean that more “weakly defended” stickleback are still able to persist in these populations, which would be reflected in reduced armour traits of fish coexisting with only SMB. Additionally, lakes with just SC show similarities to lakes with neither non-native species, similar to darter activity remaining the same with crayfish presence relative to control conditions with neither bass nor crayfish [42].

In a natural environment, if bass consume more stickleback prey when there are lower densities of SC [43], it is likely that there are other mechanisms at play driving these highly armoured phenotypes. Most of the highly armoured stickleback in the dataset were from Beaver Lake, which is very distinct and drove most of the differences between SMB + SC and other lake types. Beaver Lake has a complicated history with SMB presence (Table 2) as they were initially identified in surveys but disappeared from reports for an extended period of time. Drawing from temporal analyses, we can observe that stickleback taken from Beaver Lake in 1973 (when bass were still detected in surveys) had very long spines and numerous lateral plates, and that these values declined during the period of SMB absence. While this might suggest that stickleback armour is fluctuating with bass presence, Trout Lake also saw similar declines and has consistently contained only cutthroat trout. Prickly sculpin were also observed in Beaver Lake, which are known intraguild predators of stickleback and are suspected of having an influence on stickleback morphology [44]. Accordingly, sculpin may be partially to fully responsible for Beaver Lake stickleback armour trait values that have been considerably larger compared to other populations since at least 1973.

The temporal datasets highlight several other notable differences in stickleback traits among lakes. For the most part, both datasets A and B agree with each other, which suggests that the measurements of museum specimens by two different individuals are consistent enough to illustrate general trends. Both Klein and Dougan Lake fish have a fairly large increase in size traits for both datasets. Although present-day fish from Dougan and Klein Lakes are not distinctly larger than all the other lakes (Cusheon Lake fish tend be largest), they do appear to be clustered separately from lakes without SC or SMB. Because both of these lakes contain crayfish, it is possible that long-term exposure to SC is leading to relatively larger fish in these lakes, although it is unclear whether this transition occurred throughout the 43-year period or over a shorter time frame immediately following crayfish establishment. Although this suggests that crayfish presence favours a more benthic-like phenotype in these populations, we note that most contemporary populations resembled the benthic form in Enos Lake.

Morphological differences among lakes and non-native species combinations extend to body shape in a similar way. SMB + SC lakes are very distinct from SMB lakes, as previously described in the trait analysis. However, there are also clear shape differences between SMB and SC lake fish, whereas lakes with neither non-native species have body shapes that are midway between all lake types. The largest amount of separation occurs in PC1, where SMB lake fish have a short ascending branch of the pelvic girdle and slightly longer abdomen. SMB + SC lakes and SC lakes have stickleback with a fairly long ascending branch and a rounder head. This extended pelvic girdle shape, which is positively associated with other defensive traits such as pelvic spines and lateral plates [64], agrees with the trait analyses that identified greater armour in SMB + SC lakes and less armour in SMB lakes. Although SC lake fish have armour traits that are not especially distinct, pelvic girdle shape seems to deviate from that trend. With PC2, patterns are slightly less clear, although SMB + SC lakes appear to have fish with marginally longer and thinner tails and longer dorsal and anal fins. Previous models have predicted that longer fins positioned more caudally along the tail allows for quicker evasion in habitats occupied by other predatory fish [17]. If selection is favouring reduced interactions with both bass and crayfish, it is possible that this would select for tails more suited for evasion and speed.

The temporal analysis of body shape suggests that Beaver Lake may have been more similar to other populations in the past. Between 1973 and 2016, Beaver Lake individuals have developed rounder and larger heads, whereas there was little change in the ascending pelvic branch. Conversely, the other three lakes all had flatter backs and longer dorsal and anal fins. The ascending pelvic branch increased in Klein Lake fish, in agreement with the observation that contemporary Klein Lake fish have some of the largest ascending branches. With 2016 fish, Klein and Beaver Lake stickleback are on the upper end of PC1 and appear to have transitioned towards that extreme over time, whereas Dougan and Trout Lake fish are more intermediate along PC1 and have followed similar changes in shape over time. This reinforces that these two SC lakes (i.e., Klein and Dougan) vary not only in shape differences but with changes in shape over time, and that signal crayfish may not be the primary force driving variation in body shape in these lakes.

Interactions between SMB and SC are complex, and there is evidence here that sharing an ecosystem plays a role in their impact on threespine stickleback. Bass and crayfish are known to have a predator-prey relationship even in their native ranges, with bass also able to alter crayfish behavior and habitat use [65]. Although bass are capable of preying primarily on crayfish when they coexist in small lakes, bass will sometimes switch to diets consisting mostly of prey fish [43]. When SC coexist with bass or other predatory fish, they actively seek refuges more frequently and can outcompete native species for these shelters, including other native crayfish [66]. Predator-prey interactions can also be modified by manipulation of microhabitat structure in other systems. For example, largemouth bass growth rates and numbers increase with the removal of vegetation to create additional edges, allowing for more predation opportunities on prey that rely on vegetation for shelter [67]. Availability of refuges in the lakes sampled in this study could determine SMB foraging opportunities and interactions with stickleback and SC prey. As previously mentioned, competition with SC for shelter paired with predation by SMB could be an explanation for the increased armour and greater sizes of stickleback from the two SMB + SC lakes.

Beyond establishing statistical significance of trait and shape differences among lakes and over time, it is important to consider these differences in a biological context. The significant differences in armour traits among lakes were characterized in part by dorsal and pelvic spine lengths. Stickleback spines are an effective means of deterring SMB or other piscivores by increasing stickleback diameter and by damaging mouth parts [68]. In addition, spines have a demonstrated ability to increase the difficulty of consumption and handling time when swallowed tail first [69,70]. Evasion from trout predation is more likely as the cross-sectional diameter of stickleback approaches the predator’s mouth diameter [70]. Accordingly, body depth, pelvic girdle width, and spine lengths collectively aid in defense against gape-limited piscivores. Interpreting the biological significance of change in other traits, such as lateral plates, may be less straightforward. Fish with more lateral plates are better defended against larger fish, in part because they provide structural support for spines [71]. Conversely, individuals with more plates have lower swimming speeds, which may hamper evasion [72].

Whereas stickleback spines and plates can protect from gape-limited fish, the same might not be true for smaller invertebrate predators, including crayfish. The relationship between invertebrate predation and stickleback morphology is less certain. Dragonfly nymphs and giant water bugs showed no preference for presence or absence of spines and pelvic structures in brook stickleback, although nymphs tended to prefer smaller size classes [73]. Threespine stickleback in controlled environments with invertebrate predators resulted in greater selection for reduced armour traits and larger juveniles [27,74]. Signal crayfish have been observed preying on stickleback in the Vancouver Island region, but it is unclear to what extent this occurs or what techniques they use to catch stickleback prey [32]. Although signal crayfish may feed primarily on detritus and invertebrates [75], there is experimental evidence for their ability to prey on fish [76,77]. We expect they would most successfully prey on smaller individuals, which may contribute to the larger body sizes observed over time in Klein and Dougan Lakes.

An interesting alternative hypothesis is that their effects on stickleback reproduction, habitat, and behaviour may be driving these differences. Increased aggressive behaviour of limnetic males toward signal crayfish was observed in Enos Lake, with the suggestion that fewer limnetic males available for breeding could have driven hybridization toward more benthic-like individuals [33]. If similar behaviour is exhibited by thinner, smaller, and more aggressive individuals in non-species pair populations of stickleback, stockier, larger, and more benthic-like individuals may have greater reproductive success and, thus, would become more predominant in the population. Given uncertainty regarding the impacts of these two exotic species, together with the results presented here, we hope our study will spur additional monitoring of these and other populations, along with experimental investigations, with the goal of ensuring the long-term viability of solitary stickleback populations in the region and elsewhere.

## Supplementary Materials

Supplementary Methods, Tables S1-S8, and Figures S1-S9 can be found in the accompanying Supplementary Materials document.

## Author Contributions

Conceptualization: H.M.K. and S.M.V.; fieldwork: H.M.K.; measurements: H.M.K. and S.M.V.; genetic determination of sex: H.M.K.; data curation: H.M.K. and S.M.V.; analyses: H.M.K.; writing: H.M.K. and S.M.V.; funding acquisition: H.M.K. and S.M.V.

## Institutional Review Board Statement

All procedures were carried out in accordance with Canadian Council of Animal Care standards (University of Calgary AC15-0033).

## Data Availability Statement

Data files and analysis scripts will be uploaded to Dryad after the manuscript has been accepted.

## Acknowledgements

Heather Jamniczky and Leland Jackson provided helpful feedback on the study. We are indebted to Eric Taylor and the Beaty Biodiversity Museum for their loan of historical threespine stickleback samples. We are grateful for landowners who granted access to lakes on private property, especially those at Beaver Lake Resort and Bullock Lake Farm.

