## Supplemental Methods, Tables, and Figures for "Characterizing the impacts of exotic species on the morphology of solitary threespine stickleback (*Gasterosteus aculeatus*) populations in southwestern British Columbia"

Supplemental Material Contents

**Supplemental Methods: calculating estimated stickleback diameters**

**Table S1.** Principal component analysis loadings for measurements on contemporary data.

**Table S2.** DFA classifications of contemporary data using linear trait measurements, for 10 sampled lakes and four non-native species combinations.

**Table S3.** Mean (with standard deviation and coefficient of variation) estimated diameter for each population for different times and time period.

**Table S4.** Principal component analysis for temporal datasets A and B. Trait values shown are character loadings.

**Table S5.** MANOVA results for effects of year and sex (dataset A) and year alone (dataset B) on stickleback morphology. Factors are bolded where significant i.e., when $p < 0.05$.

**Table S6.** Correlations between principal components and the first linear discriminant.

**Table S7.** Procrustes ANOVA results for the effects of year, lake, and non-native species combination on body shape. The temporal analysis is split by lake, whereas the spatial analysis is split between males and females.

**Table S8.** DFA classifications of lakes and non-native species combinations for contemporary body shape data. Randomized data consists of the original dataset with lakes randomized and attributed to a random individual's landmark data.

**Figure S1:** Landmarks (top) and measurements (middle and bottom) taken from the ventral and left lateral sides of each individual.

**Figure S2.** Proportion of correct DFA classifications for each lake and species combination, based on linear trait measurements of males and females.

**Figure S3.** Trait variation in temporal data for datasets A (top panels) and B (bottom panels).

**Figure S4.** Changes in estimated cross-sectional diameter of stickleback from 1973 to 2016, for Beaver, Dougan, Klein, and Trout Lake.

**Figure S5.** Male body shape variation among lakes and non-native species combinations.

**Figure S6.** Female body shape variation among lakes and non-native species combinations.

**Figure S7.** Proportion of correct DFA classifications for each lake and non-native species combination, based on body shape of males and females.

**Figure S8.** Body shape variation between years for Beaver and Klein Lake, explained using the first two components of a PCA on bending-corrected landmark data.

**Figure S9.** Body shape variation between years for Dougan and Trout Lake, explained using the first two components of a PCA on bending-corrected landmark data.

**Supplemental Methods: calculating estimated stickleback diameters**

Body depth, second dorsal spine length, pelvic spine length, and pelvic girdle width were used to approximate the diameter of stickleback in a defensive position. Because pelvic spines protrude at an angle, the vertical extension of these spines was calculated by assuming that the spines are at a 20.9° angle from the horizontal (following Reimchen 1991). With this angle and the length of the spine, vertical extension of the spine could be calculated. This distance, in addition to the dorsal spine and body depth lengths, forms a bisecting line in a triangle that has a circumcircle describing the cross-sectional diameter of stickleback.

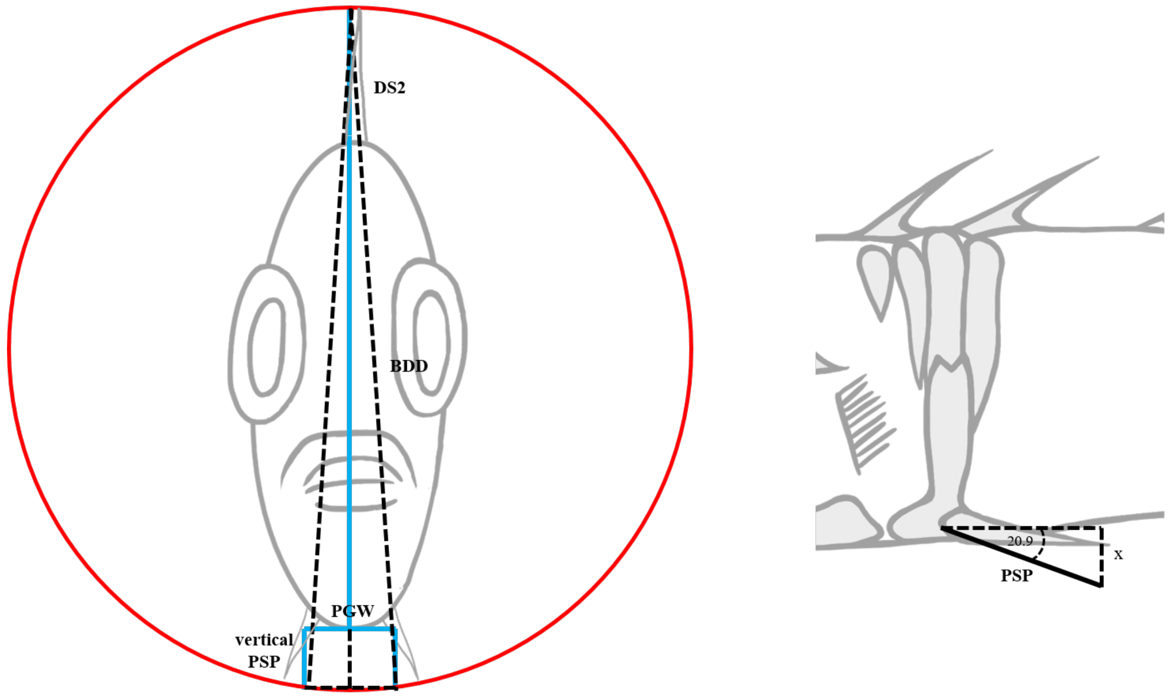


Reimchen, T. E. (1991). Trout foraging failures and the evolution of body size in stickleback. *Copeia, 1991*, 1098-1104.

**Table S1.** Principal component analysis loadings for measurements on contemporary data.

| | Female | | | Male | | |
|---|---|---|---|---|---|---|
| Trait | PC1 | PC2 | PC3 | PC1 | PC2 | PC3 |
| Standard length | 0.95 | 0.23 | 0.00 | 0.94 | 0.21 | 0.00 |
| Eye diameter | 0.90 | 0.05 | 0.04 | 0.80 | -0.01 | 0.13 |
| Gape length | 0.71 | -0.09 | 0.17 | 0.70 | -0.17 | 0.37 |
| Snout length | 0.85 | 0.14 | 0.23 | 0.74 | 0.28 | 0.44 |
| Head length | 0.96 | 0.07 | 0.11 | 0.93 | 0.10 | 0.25 |
| Head depth | 0.93 | 0.12 | -0.01 | 0.91 | 0.16 | -0.01 |
| 1st dorsal spine | 0.10 | 0.91 | 0.01 | -0.01 | 0.91 | -0.06 |
| 2nd dorsal spine | 0.10 | 0.89 | -0.06 | 0.08 | 0.92 | -0.13 |
| Body depth | 0.92 | 0.18 | -0.08 | 0.88 | 0.22 | -0.20 |
| Ectocoracoid | 0.80 | 0.31 | -0.11 | 0.62 | 0.44 | -0.10 |
| Caudal peduncle length | 0.69 | 0.09 | -0.22 | 0.63 | -0.20 | -0.33 |
| Caudal peduncle width | 0.88 | 0.19 | 0.04 | 0.83 | 0.06 | -0.08 |
| Pelvic girdle width | 0.83 | 0.11 | -0.16 | 0.65 | 0.22 | -0.40 |
| Pelvic girdle length | 0.51 | 0.68 | -0.13 | 0.33 | 0.80 | -0.13 |
| Lateral plates | -0.02 | 0.67 | 0.15 | 0.04 | 0.67 | 0.20 |
| Pelvic spine length | 0.19 | 0.91 | -0.20 | 0.08 | 0.94 | -0.10 |
| Gill rakers | 0.04 | -0.05 | 0.92 | 0.02 | -0.09 | 0.70 |
| Proportion of variance explained | 0.50 | 0.21 | 0.07 | 0.56 | 0.33 | 0.10 |

**Table S2.** DFA classifications of contemporary data using linear trait measurements, for 10 sampled lakes and four non-native species combinations.

| | | Proportion of correct classifications | | |
|---|---|---|---|---|
| | | Original dataset | Randomized dataset | Prior probabilities |
| Males | Beaver | 1.000 | 0.280 | 0.152 |
| | Blackburn | 0.400 | 0.000 | 0.030 |
| | Bullock | 0.833 | 0.778 | 0.220 |
| | Cusheon | 0.600 | 0.000 | 0.030 |
| | Dougan | 0.000 | 0.000 | 0.055 |
| | Klein | 0.600 | 0.000 | 0.091 |
| | Maple | 0.789 | 0.000 | 0.116 |
| | Stowell | 0.000 | 0.000 | 0.079 |
| | Trout | 0.667 | 0.000 | 0.110 |
| | Weston | 0.316 | 0.000 | 0.116 |
| | Overall | 0.622 | 0.213 | |
| Females | Beaver | 1.000 | 0.000 | 0.092 |
| | Blackburn | 0.633 | 0.767 | 0.196 |
| | Bullock | 0.500 | 0.000 | 0.026 |
| | Cusheon | 0.846 | 0.000 | 0.085 |
| | Dougan | 0.636 | 0.136 | 0.144 |
| | Klein | 0.850 | 0.000 | 0.131 |
| | Maple | 0.000 | 0.000 | 0.039 |
| | Stowell | 0.706 | 0.000 | 0.111 |
| | Trout | 0.333 | 0.000 | 0.078 |
| | Weston | 0.267 | 0.333 | 0.098 |
| | Overall | 0.634 | 0.183 | |
| Males | None | 0.600 | 0.680 | 0.305 |
| | SC | 0.721 | 0.000 | 0.262 |
| | SMB | 0.756 | 0.220 | 0.250 |
| | SC + SMB | 0.867 | 0.000 | 0.183 |
| | Overall | 0.720 | 0.262 | |
| Females | None | 0.500 | 0.523 | 0.288 |
| | SC | 0.771 | 0.396 | 0.314 |
| | SMB | 0.618 | 0.206 | 0.222 |
| | SC + SMB | 0.593 | 0.327 | 0.176 |
| | Overall | 0.627 | 0.327 | |

**Table S3.** Mean (with standard deviation and coefficient of variation) estimated diameter for each population for different times and time period.

| | Lake | Year | Mean (mm) | SD (mm) | CV (%) |
|---|---|---|---|---|---|
| Contemporary analysis | BB | 2016 | 14.637 | 1.715 | 11.719 |
| | BU | 2016 | 14.916 | 1.254 | 8.410 |
| | BV | 2016 | 19.818 | 2.009 | 10.135 |
| | CU | 2016 | 21.476 | 1.545 | 7.193 |
| | DG | 2016 | 18.675 | 2.005 | 10.737 |
| | KL | 2016 | 16.774 | 1.332 | 7.941 |
| | MP | 2016 | 18.044 | 1.347 | 7.466 |
| | SW | 2016 | 15.908 | 1.407 | 8.842 |
| | TR | 2016 | 15.527 | 1.282 | 8.254 |
| | WE | 2016 | 15.794 | 1.440 | 9.118 |
| Temporal analysis | BV | 1970-1982 | 19.785 | 2.251 | 11.380 |
| | | 2016 | 19.818 | 2.009 | 10.135 |
| | DG | 1970-1982 | 14.731 | 1.866 | 12.670 |
| | | 2016 | 18.675 | 2.005 | 10.737 |
| | KL | 1970-1982 | 14.600 | 1.644 | 11.257 |
| | | 2016 | 16.774 | 1.332 | 7.941 |
| | TR | 1970-1982 | 15.925 | 1.778 | 11.164 |
| | | 2016 | 15.527 | 1.282 | 8.254 |

**Table S4.** Principal component analysis for temporal datasets A and B. Trait values shown are character loadings.

| Dataset | Traits | PC1 | PC2 |
|---|---|---|---|
| A | Standard length | 0.13 | 0.96 |
| | Body depth | 0.19 | 0.96 |
| | 1st dorsal spine | 0.93 | 0.20 |
| | 2nd dorsal spine | 0.91 | 0.27 |
| | Lateral plates | 0.76 | -0.06 |
| | Pelvic spine length | 0.92 | 0.28 |
| | **Proportion of variance explained** | **0.53** | **0.34** |
| B | Standard length | 0.92 | 0.26 |
| | Eye diameter | 0.85 | 0.12 |
| | Gape length | 0.71 | 0.02 |
| | Snout length | 0.44 | 0.36 |
| | Head length | 0.95 | 0.17 |
| | Head depth | 0.94 | 0.15 |
| | 1st dorsal spine | 0.13 | 0.87 |
| | 2nd dorsal spine | 0.12 | 0.90 |
| | Body depth | 0.92 | 0.19 |
| | Ectocoracoid | 0.65 | 0.53 |
| | Caudal peduncle length | 0.50 | 0.23 |
| | Caudal peduncle width | 0.90 | 0.11 |
| | Pelvic girdle width | 0.70 | 0.20 |
| | Pelvic girdle length | 0.26 | 0.84 |
| | Lateral plates | 0.12 | 0.59 |
| | Pelvic spine length | 0.11 | 0.94 |
| | **Proportion of variance explained** | **0.44** | **0.26** |

**Table S5.** MANOVA results for effects of year and sex (dataset A) and year alone (dataset B) on stickleback morphology. Factors are bolded where significant i.e., when p < 0.05.

| Dataset | Lake | | df | Pillai | F | num df | den df | p value |
|---|---|---|---|---|---|---|---|---|
| A | Bullock | **Year** | 1 | 0.245 | 4.212 | 2 | 26 | 0.026 |
| | | Sex | 1 | 0.012 | 0.155 | 2 | 26 | 0.857 |
| | | Year x Sex | 1 | 0.118 | 1.735 | 2 | 26 | 0.197 |
| | | Residuals | 27 | | | | | |
| | Beaver | **Year** | 1 | 0.273 | 13.734 | 2 | 73 | < 0.001 |
| | | **Sex** | 1 | 0.157 | 6.779 | 2 | 73 | 0.002 |
| | | **Year x Sex** | 1 | 0.235 | 11.224 | 2 | 73 | < 0.001 |
| | | Residuals | 74 | | | | | |
| | Dougan | **Year** | 1 | 0.526 | 27.709 | 2 | 50 | < 0.001 |
| | | **Sex** | 1 | 0.144 | 4.195 | 2 | 50 | 0.021 |
| | | Year x Sex | 1 | 0.088 | 2.414 | 2 | 50 | 0.100 |
| | | Residuals | 51 | | | | | |
| | Klein | **Year** | 1 | 0.633 | 59.466 | 2 | 69 | < 0.001 |
| | | **Sex** | 1 | 0.197 | 8.485 | 2 | 69 | < 0.001 |
| | | Year x Sex | 1 | 0.063 | 2.300 | 2 | 69 | 0.108 |
| | | Residuals | 70 | | | | | |
| | Trout | **Year** | 1 | 0.121 | 3.438 | 2 | 50 | 0.040 |
| | | Sex | 1 | 0.026 | 0.671 | 2 | 50 | 0.516 |
| | | Year x Sex | 1 | 0.003 | 0.072 | 2 | 50 | 0.930 |
| | | Residuals | 51 | | | | | |
| B | Beaver | **Year** | 1 | 0.682 | 82.611 | 2 | 77 | < 0.001 |
| | | Residuals | 78 | | | | | |
| | Dougan | **Year** | 1 | 0.489 | 31.623 | 2 | 66 | < 0.001 |
| | | Residuals | 67 | | | | | |
| | Klein | **Year** | 1 | 0.702 | 67.179 | 2 | 57 | < 0.001 |
| | | Residuals | 58 | | | | | |
| | Trout | **Year** | 1 | 0.658 | 61.702 | 2 | 64 | < 0.001 |
| | | Residuals | 65 | | | | | |

**Table S6.** Correlations between principal components and the first linear discriminant.

| | | Correlation with linear discriminant 1 | |
|---|---|---|---|
| | | PC1 | PC2 |
| Spatial analysis | Lake | -0.208 | 0.978 |
| | Species combination | 0.483 | 0.876 |
| Temporal analysis (A) | Bullock | 0.016 | -0.978 |
| | Beaver | -0.973 | -0.257 |
| | Dougan | 0.655 | 0.770 |
| | Klein | 0.310 | 0.998 |
| | Trout | -0.971 | 0.347 |
| Temporal analysis (B) | Beaver | 0.501 | -0.863 |
| | Dougan | 0.996 | 0.456 |
| | Klein | 0.950 | -0.381 |
| | Trout | 0.276 | -0.932 |

**Table S7.** Procrustes ANOVA results for the effects of year, lake, and non-native species combination on body shape. The temporal analysis is split by lake, whereas the spatial analysis is split between males and females.

| | | | Df | SS | MS | Rsq | F | Z | p |
|---|---|---|---|---|---|---|---|---|---|
| Temporal analysis | Beaver | **Year** | 1 | 0.010 | 0.014 | 0.067 | 5.632 | 4.690 | 0.001 |
| | | Residuals | 78 | 0.144 | 0.002 | | | | |
| | | Total | 79 | 0.154 | | | | | |
| | Dougan | **Year** | 1 | 0.016 | 0.016 | 0.102 | 7.645 | 5.875 | 0.002 |
| | | Residuals | 67 | 0.143 | 0.002 | | | | |
| | | Total | 68 | 0.159 | | | | | |
| | Klein | **Year** | 1 | 0.012 | 0.012 | 0.104 | 6.646 | 5.359 | 0.001 |
| | | Residuals | 57 | 0.106 | 0.002 | | | | |
| | | Total | 58 | 0.118 | | | | | |
| | Trout | **Year** | 1 | 0.014 | 0.014 | 0.106 | 7.695 | 6.008 | 0.001 |
| | | Residuals | 65 | 0.120 | 0.002 | | | | |
| | | Total | 66 | 0.135 | | | | | |
| Spatial analysis | Males | **Lake** | 9 | 0.118 | 0.013 | 0.391 | 10.758 | 6.689 | 0.001 |
| | | Residuals | 151 | 0.185 | 0.001 | | | | |
| | | Total | 160 | 0.303 | | | | | |
| | Females | **Lake** | 9 | 0.118 | 0.013 | 0.345 | 7.963 | 5.390 | 0.001 |
| | | Residuals | 136 | 0.225 | 0.002 | | | | |
| | | Total | 145 | 0.343 | | | | | |
| | Males | **Sp. combo** | 3 | 0.072 | 0.023 | 0.237 | 16.235 | 11.454 | 0.001 |
| | | Residuals | 157 | 0.231 | 0.001 | | | | |
| | | Total | 160 | 0.303 | | | | | |
| | Females | **Sp. combo** | 3 | 0.063 | 0.021 | 0.183 | 10.635 | 8.267 | 0.001 |
| | | Residuals | 142 | 0.280 | 0.002 | | | | |
| | | Total | 145 | 0.343 | | | | | |

**Table S8.** DFA classifications of lakes and non-native species combinations for contemporary body shape data. Randomized data consists of the original dataset with lakes randomized and attributed to a random individual's landmark data.

| | | Proportion of correct classifications | | |
|---|---|---|---|---|
| | | Original dataset | Randomized data | Prior probabilities |
| Males | Blackburn | 1.000 | 0.000 | 0.031 |
| | Bullock | 1.000 | 0.457 | 0.217 |
| | Beaver | 1.000 | 0.040 | 0.155 |
| | Cusheon | 1.000 | 0.200 | 0.031 |
| | Dougan | 1.000 | 0.000 | 0.056 |
| | Klein | 1.000 | 0.067 | 0.093 |
| | Maple | 1.000 | 0.056 | 0.112 |
| | Stowell | 1.000 | 0.000 | 0.081 |
| | Trout | 1.000 | 0.000 | 0.112 |
| | Weston | 1.000 | 0.167 | 0.112 |
| | Overall | 1.000 | 0.143 | |
| Females | Blackburn | 1.000 | 0.200 | 0.171 |
| | Bullock | 1.000 | 0.000 | 0.027 |
| | Beaver | 1.000 | 0.143 | 0.096 |
| | Cusheon | 0.923 | 0.000 | 0.089 |
| | Dougan | 0.955 | 0.167 | 0.151 |
| | Klein | 1.000 | 0.211 | 0.130 |
| | Maple | 1.000 | 0.000 | 0.041 |
| | Stowell | 1.000 | 0.188 | 0.110 |
| | Trout | 1.000 | 0.000 | 0.082 |
| | Weston | 1.000 | 0.200 | 0.103 |
| | Overall | 0.986 | 0.130 | |
| Males | None | 0.878 | 0.245 | 0.304 |
| | SC | 0.952 | 0.238 | 0.261 |
| | SMB | 1.000 | 0.400 | 0.248 |
| | SC + SMB | 0.900 | 0.167 | 0.186 |
| | Overall | 0.932 | 0.267 | |
| Females | None | 0.907 | 0.349 | 0.295 |
| | SC | 0.723 | 0.234 | 0.322 |
| | SMB | 0.931 | 0.207 | 0.199 |
| | SC + SMB | 0.889 | 0.111 | 0.185 |
| | Overall | 0.849 | 0.240 | |

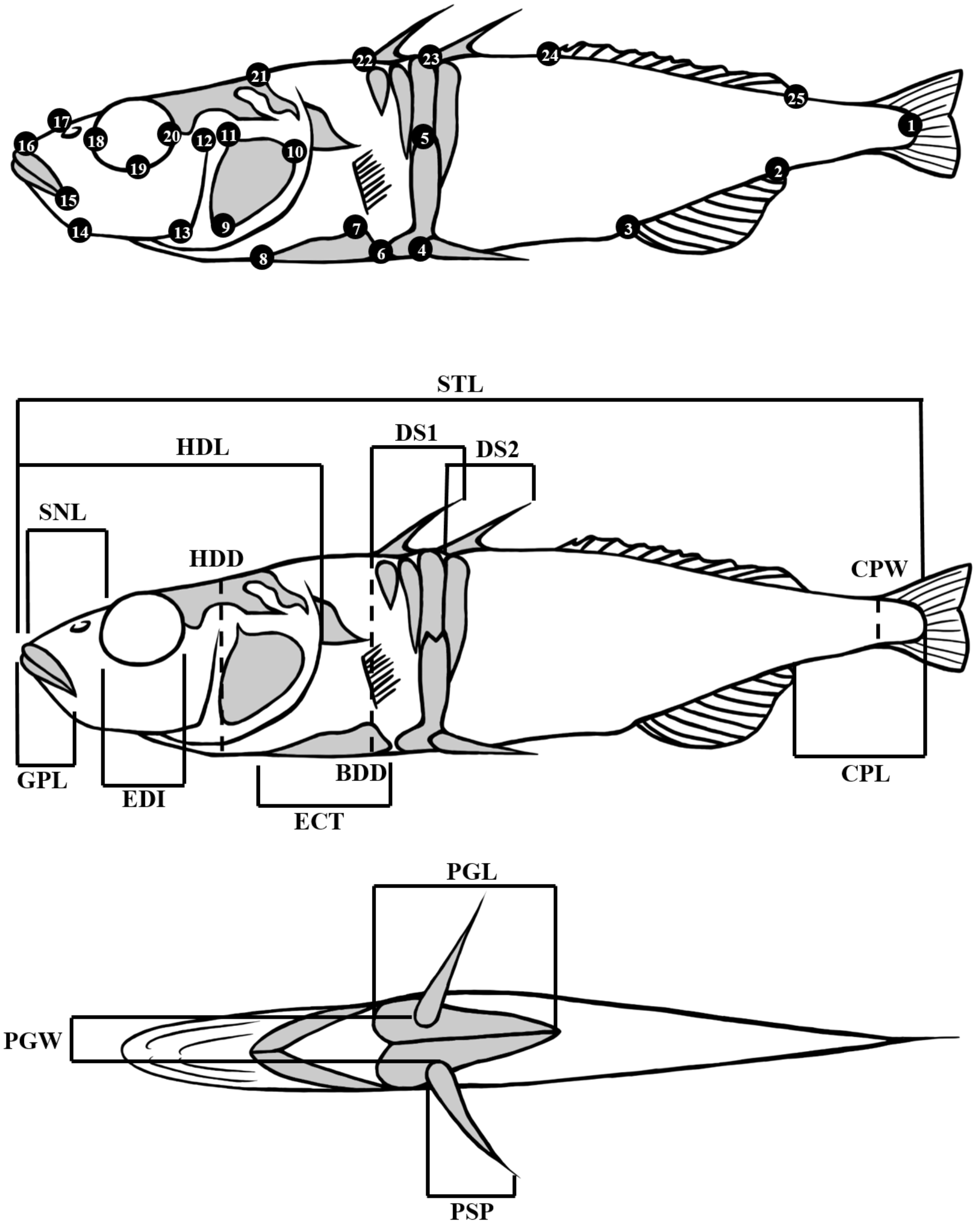


**Figure S1:** Landmarks (top) and measurements (middle and bottom) taken from the ventral and left lateral sides of each individual. Order of landmark placement is indicated by numbered circles. Measurements are indicated by abbreviations as follows: BDD = body depth, CPL = caudal peduncle length, CPW = caudal peduncle width, DS1 = 1$^{st}$ dorsal spine, DS2 = 2$^{nd}$ dorsal spine, ECT = ectocoracoid length, EDWE = eye diameter, GPL = gape length, HDL = head length, HDW = head width, PGL = pelvic girdle length, PGW = pelvic girdle width, PSP = pelvic spine length, SNL = snout length, STL = standard length.

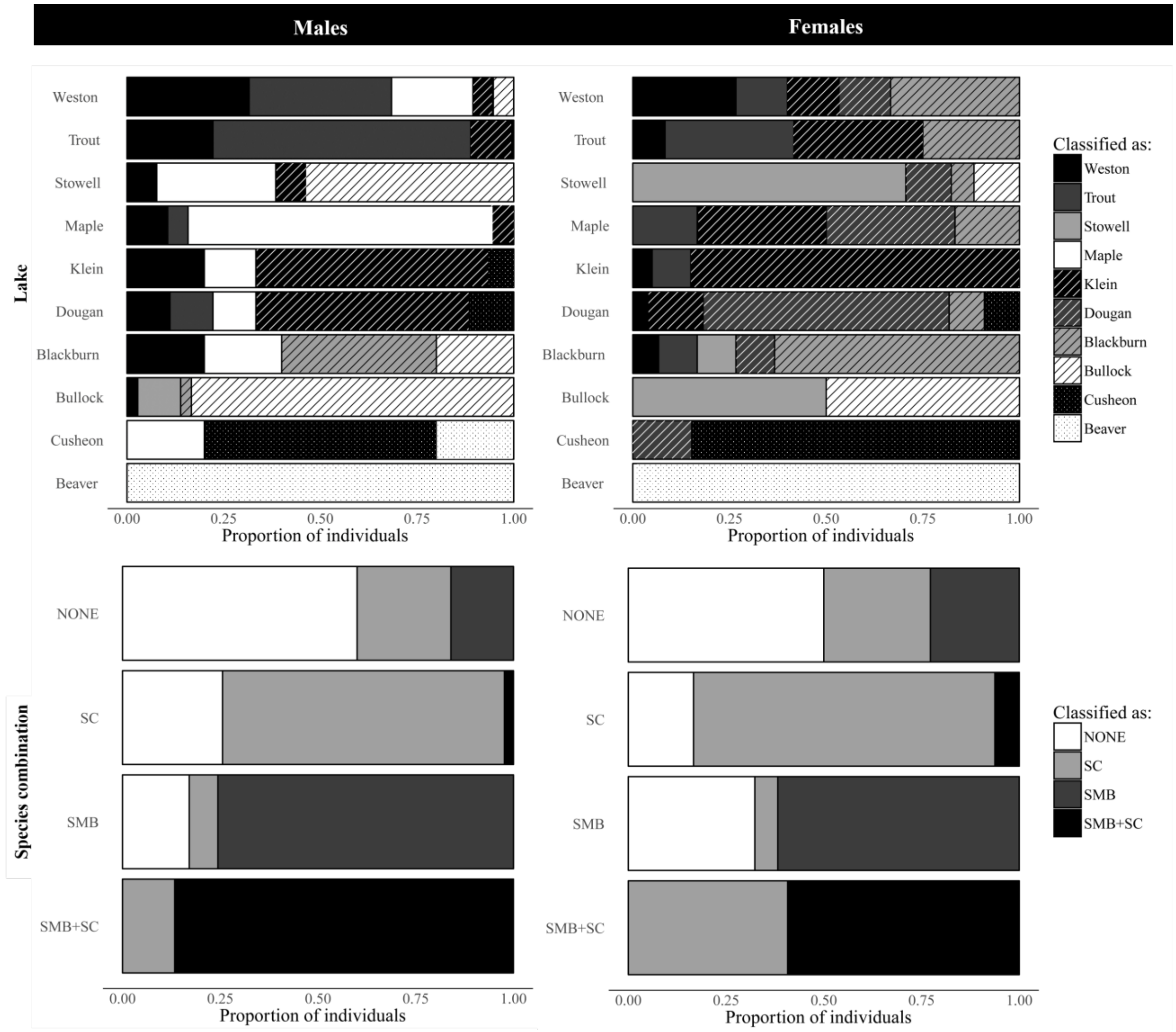


**Figure S2.** Proportion of correct DFA classifications for each lake and species combination, based on linear trait measurements of males and females. Each bar represents the total individuals from each lake or non-native species combination, and the shading and pattern segments within each bar represent the classification given to each proportion of individuals.

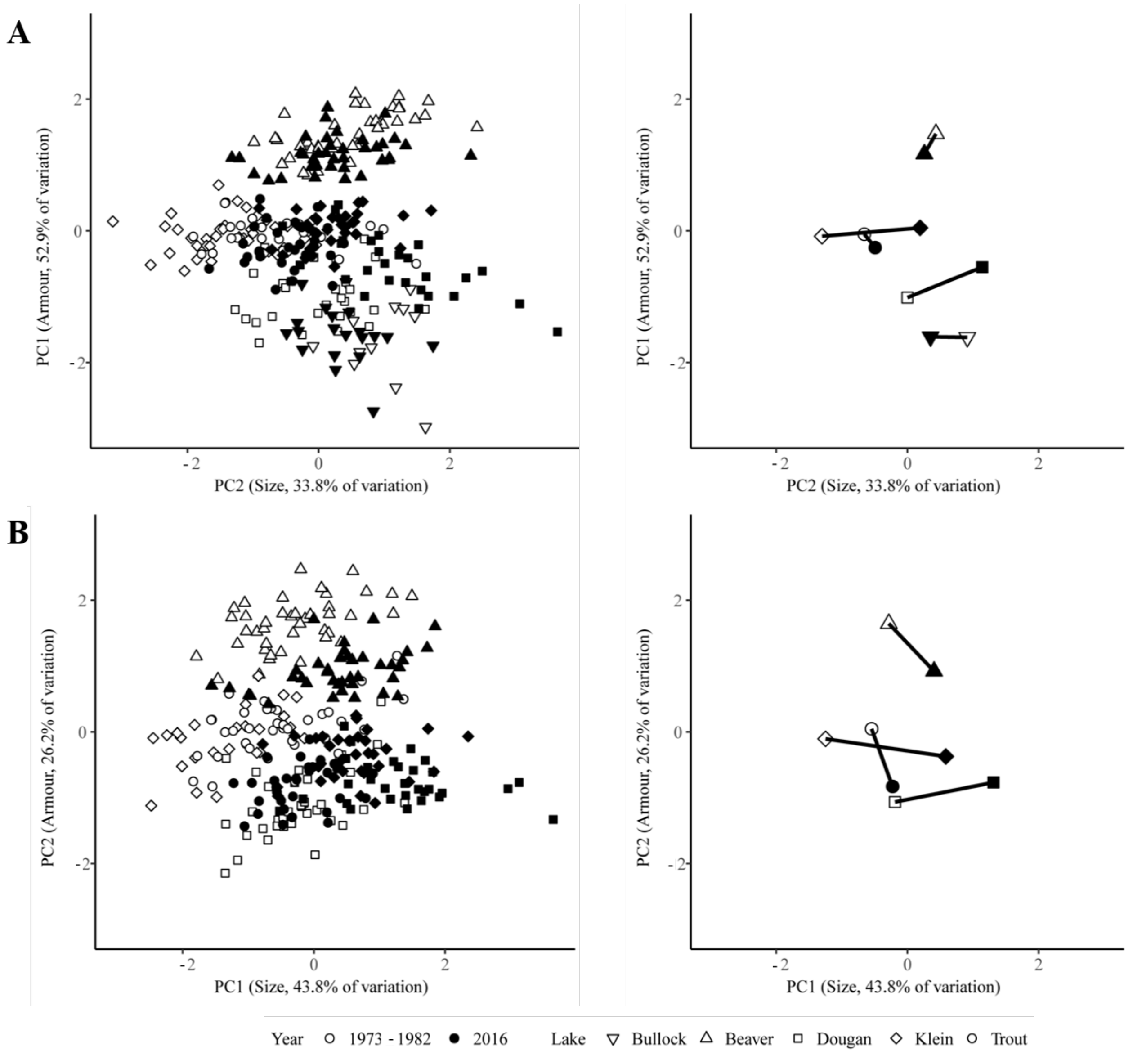


**Figure S3.** Trait variation in temporal data for datasets A (top panels) and B (bottom panels). Principal Component 1 primarily represents size-related traits whereas principal component 2 represents armour traits. The left column shows individuals whereas the right column shows mean scores for each lake and year. Lines represent the mean change from past to present.

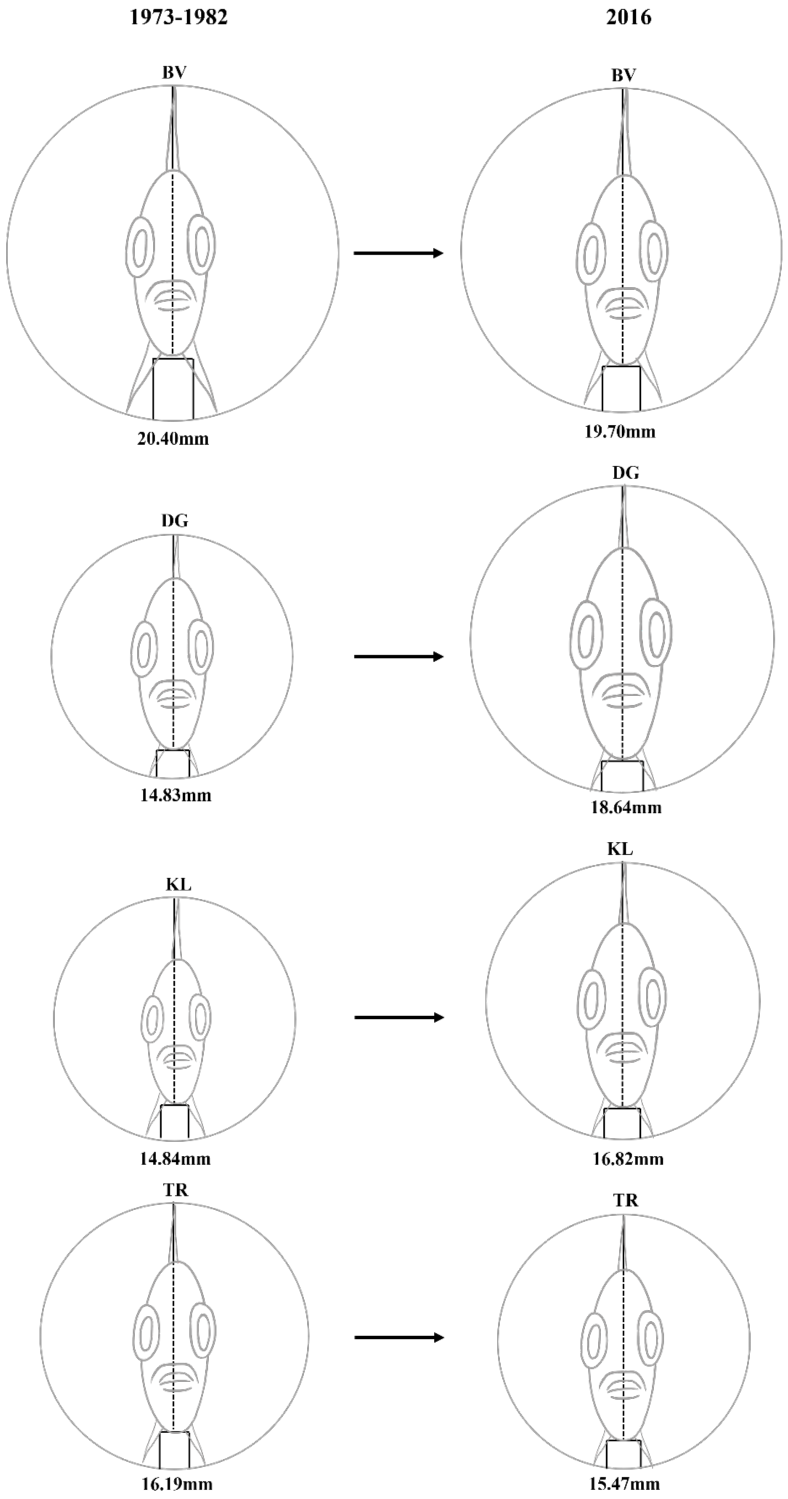


**Figure S4.** Changes in estimated cross-sectional diameter of stickleback from 1973 to 2016, for Beaver, Dougan, Klein, and Trout Lake.

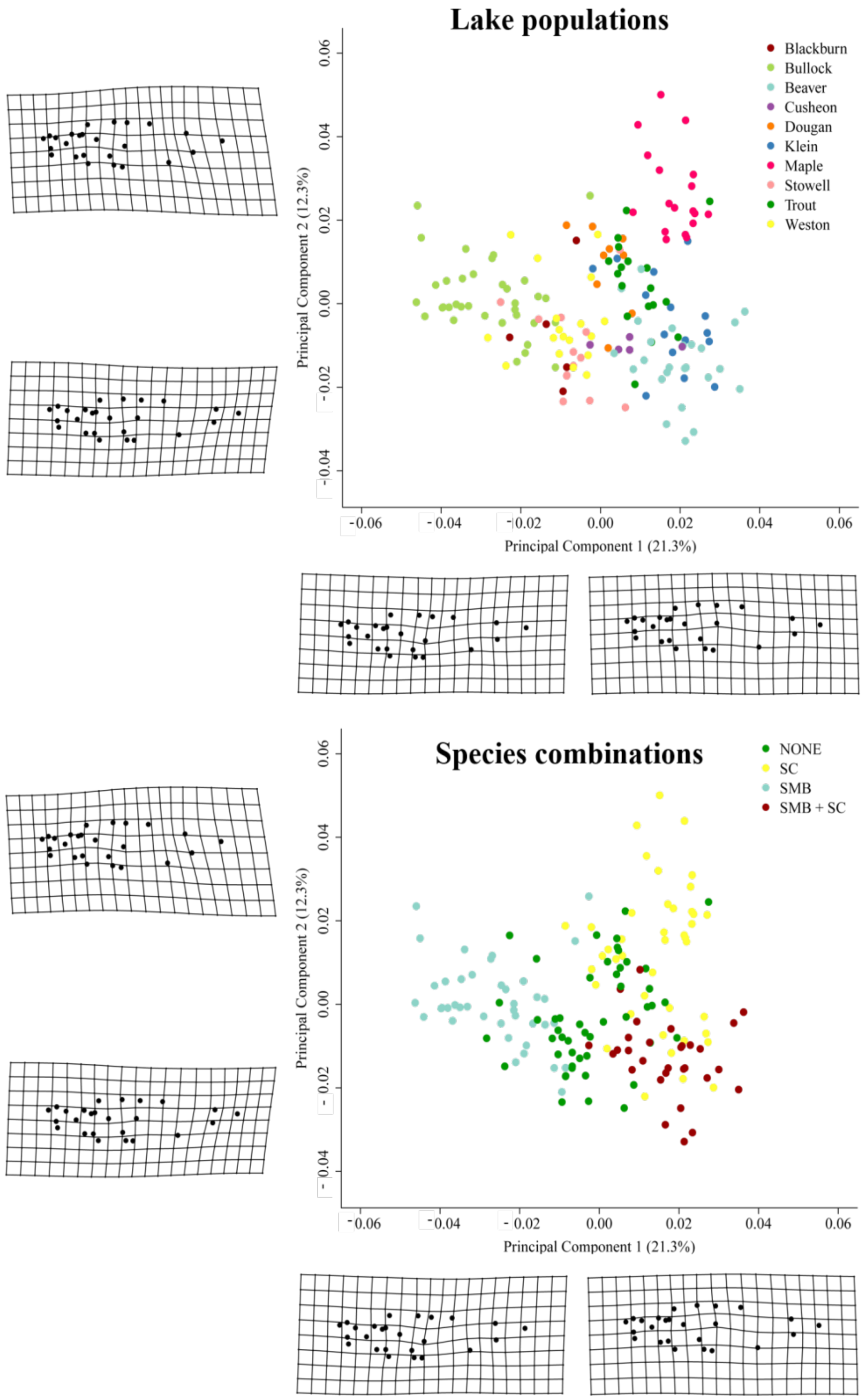


**Figure S5.** Male body shape variation among lakes and non-native species combinations, explained using the first two components of a PCA on bending-corrected landmark data. Warp meshes represent maximum and minimum shape relative to the mean, along each axis.

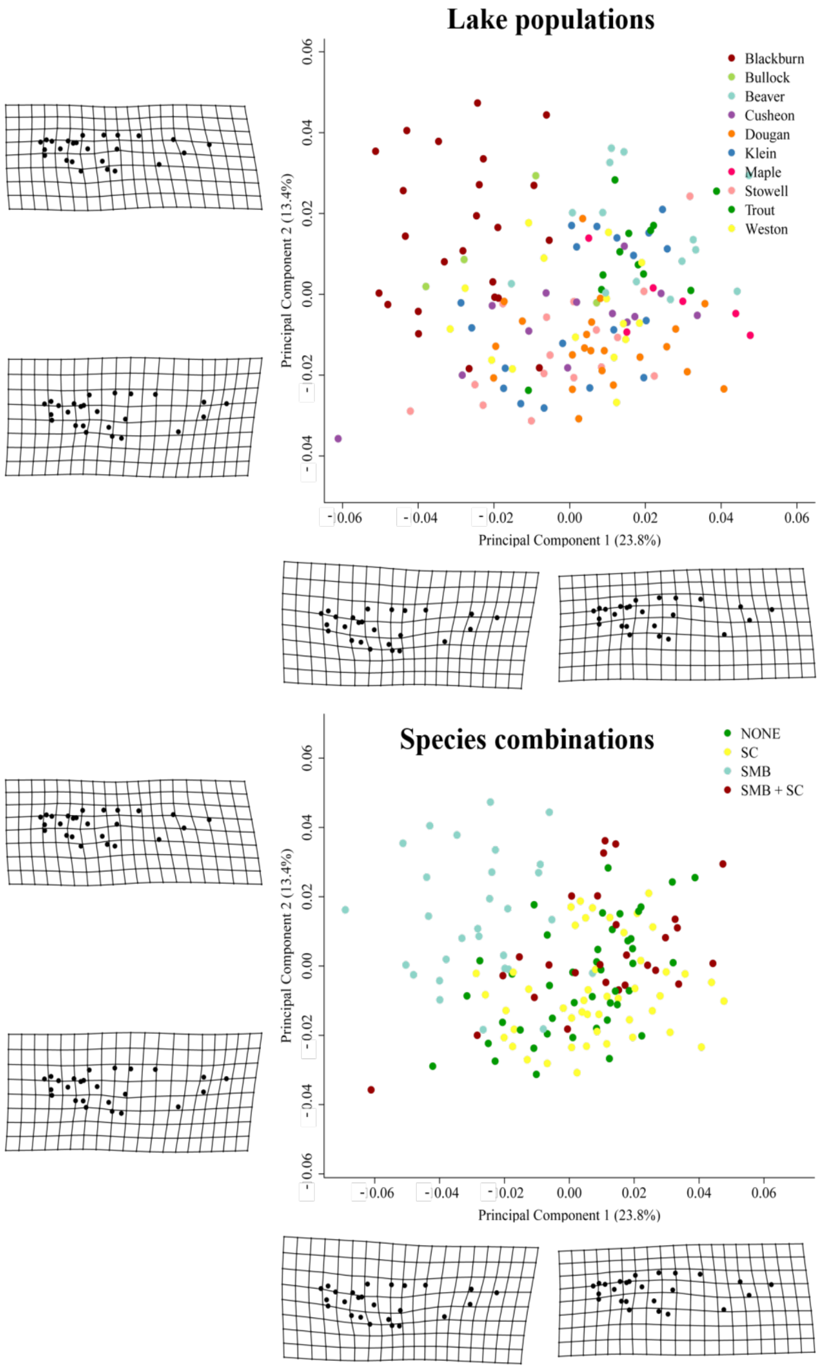


**Figure S6.** Female body shape variation among lakes and non-native species combinations, explained using the first two components of a PCA on bending-corrected landmark data. Warp meshes represent maximum and minimum shape relative to the mean, along each axis.

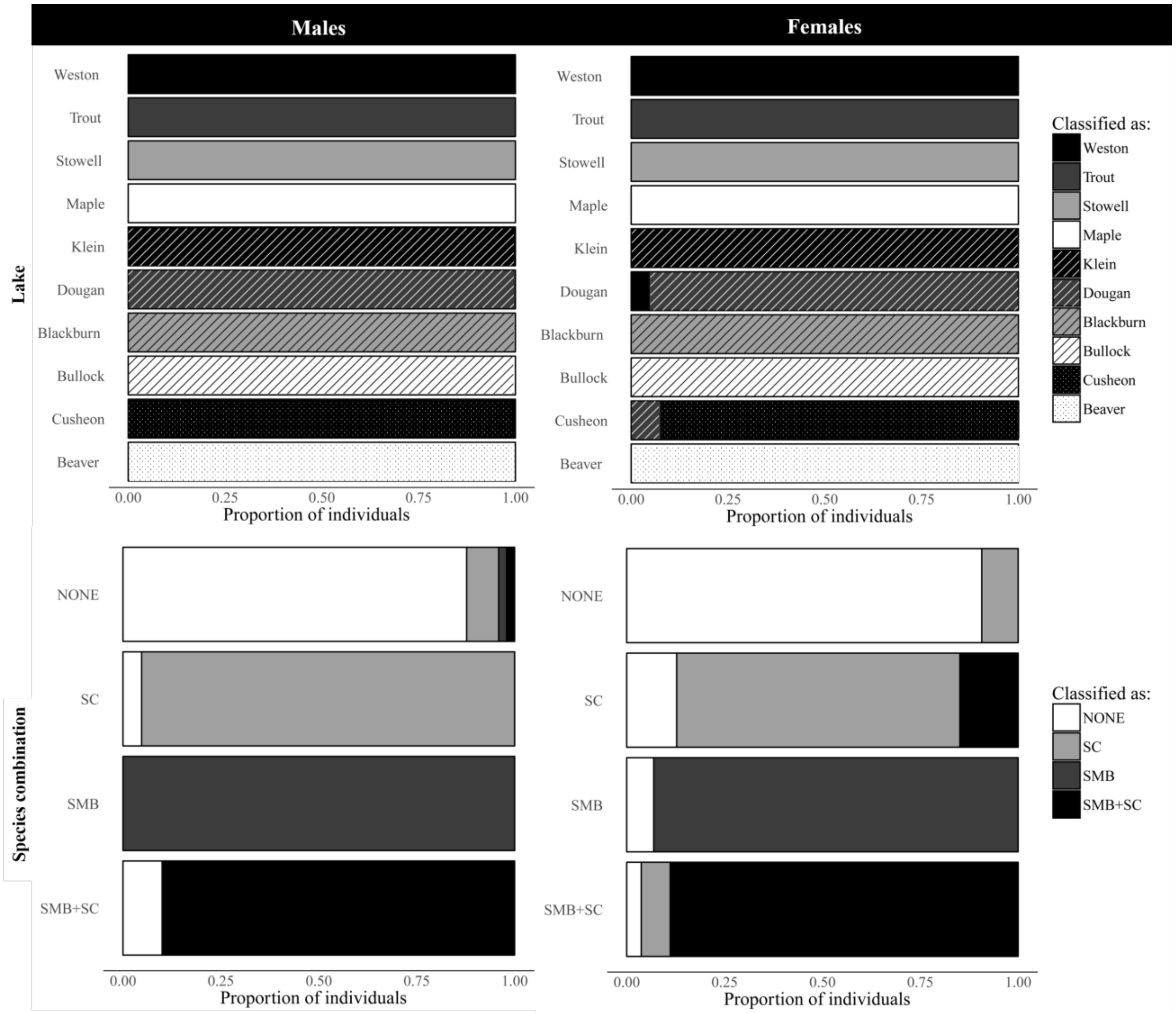


**Figure S7.** Proportion of correct DFA classifications for each lake and non-native species combination, based on body shape of males and females. Each bar represents the total individuals from each lake or non-native species combination, and the shading and pattern segments within each bar represent the classification given to each proportion of individuals.

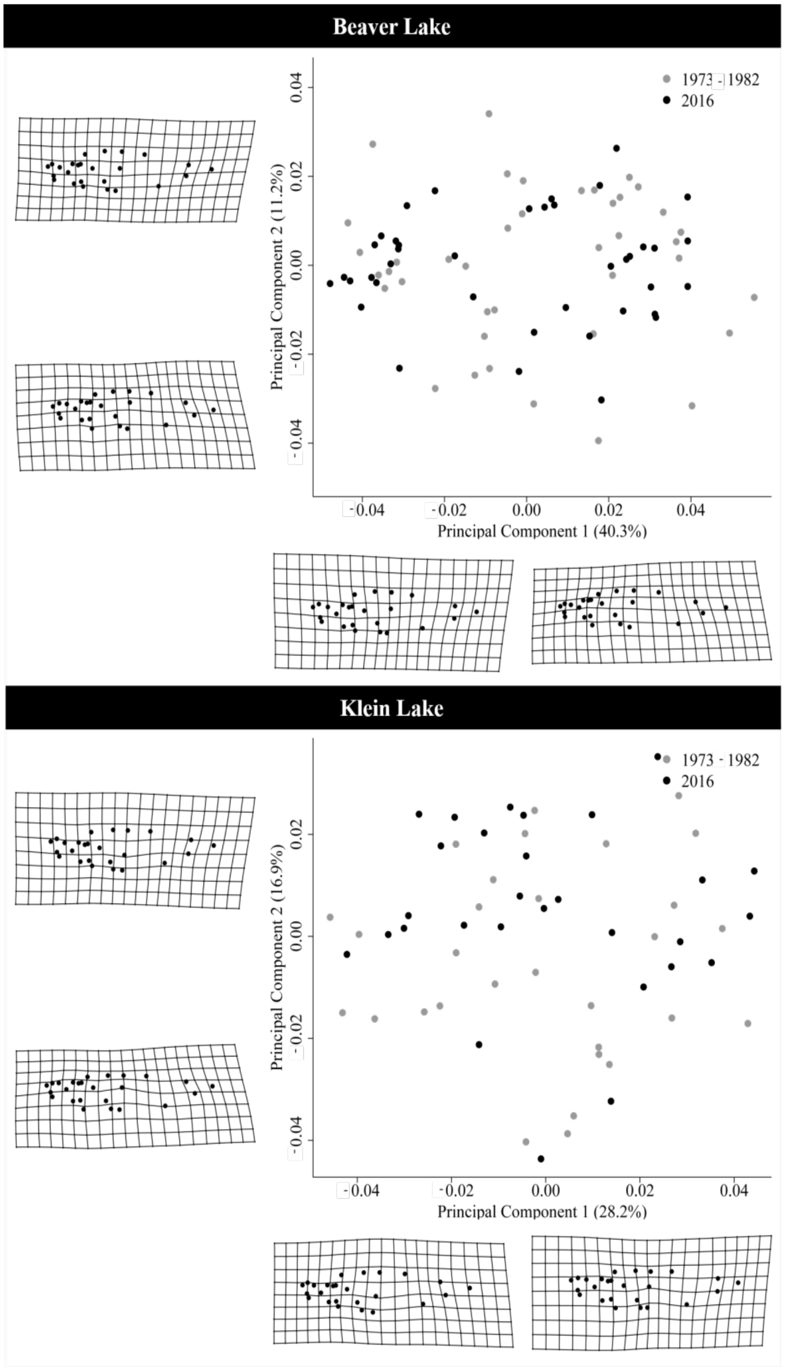


**Figure S8.** Body shape variation between years for Beaver and Klein Lake, explained using the first two components of a PCA on bending-corrected landmark data. Warp meshes represent maximum and minimum shape relative to the mean, along each axis.

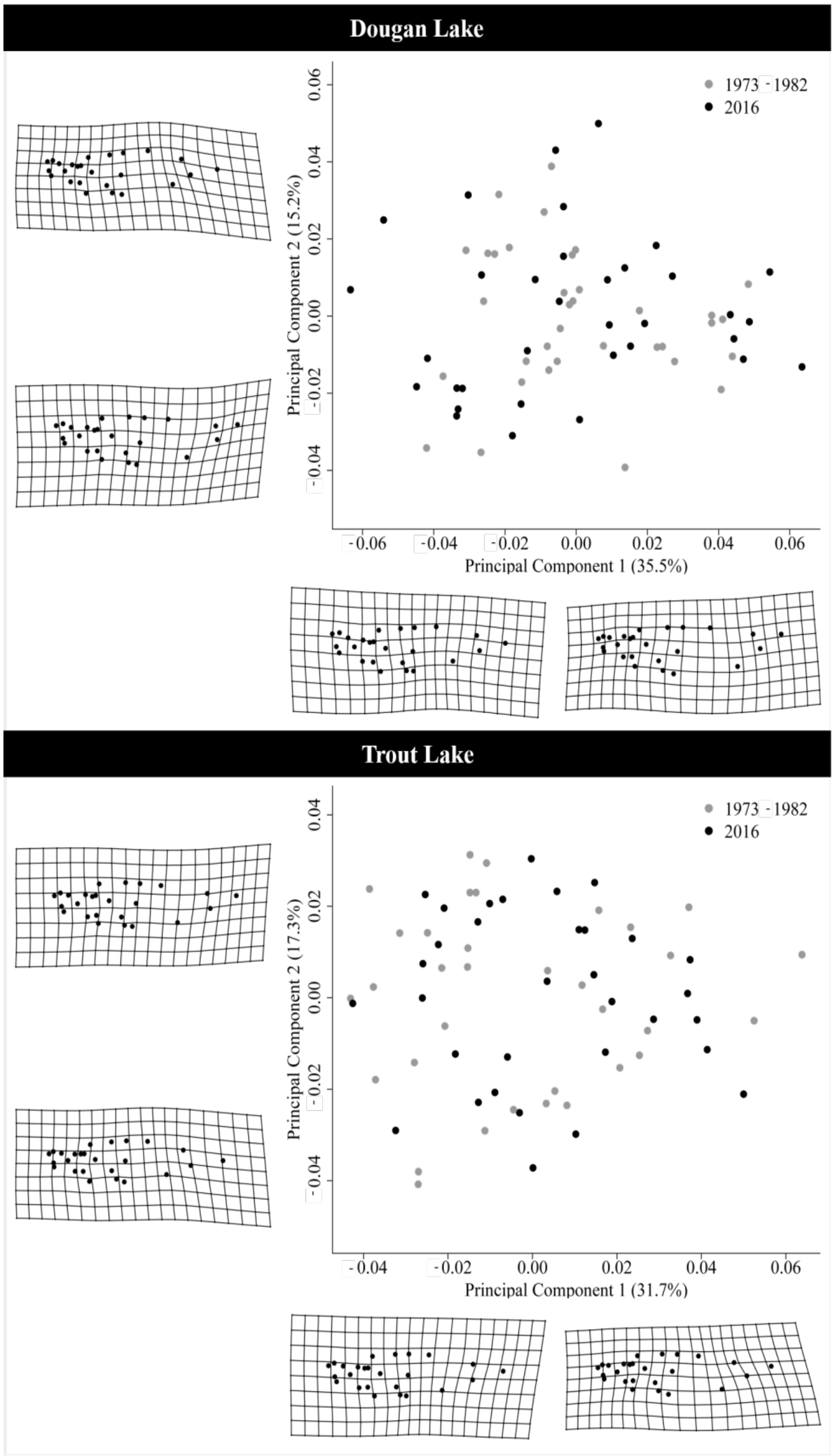


**Figure S9.** Body shape variation between years for Dougan and Trout Lake, explained using the first two components of a PCA on bending-corrected landmark data. Warp meshes represent maximum and minimum shape relative to the mean, along each axis.
